# A chromosome-scale reference genome and single-nucleus root atlas reveal the cellular and molecular basis of coastal adaptation in sand bean (*Strophostyles helvola*)

**DOI:** 10.64898/2026.08.27.747562

**Authors:** Quang Tri Le, Xuebo Zhao, Ping Yates, Henry W. Schmidt, Matias Kirst, Qiuyun Jenny Xiang, Chau Tran, Song Li, Zhangjun Fei, Bao-Hua Song

## Abstract

Plants adapted to saline coastal habitats provide valuable systems for understanding how natural selection reshapes stress-response programs, yet the cellular and molecular basis of this adaptation remains poorly resolved. Here, we generated a chromosome-scale reference genome and a single-nucleus transcriptomic atlas of roots from sand bean (*Strophostyles helvola*), a wild legume represented by salt-tolerant Beach and salt-sensitive Inland ecotypes. The atlas comprised 87,901 nuclei assigned to 25 transcriptional clusters representing 16 major root cell types. Salt exposure elicited cell-type-specific transcriptional responses that differed between Beach and Inland roots. Beach roots preferentially maintained respiration-, energy metabolism-, and protein-homeostasis-associated functions, whereas Inland roots showed stronger induction of canonical abiotic-stress, ABA, water-deficit, hypoxia, and oxidative-stress programs. Integrating baseline ecotype differences with salt-responsive expression revealed that many genes induced by salt in Inland roots were already expressed at higher levels in untreated Beach roots. This baseline-enriched configuration comprised a broadly distributed regulatory backbone, including ERF-family transcription factors and RING-type E3 ubiquitin ligases, together with cell-type-associated modules involving ion and water transport at the root-soil interface, redox and dehydration protection in outer-root tissues, endodermal barrier-associated genes, and vascular regulatory candidates. These findings support a model in which Beach salt tolerance is associated not with entirely distinct stress-response pathways, but with the pre-existing and spatially organized deployment of conserved protective programs that remain responsive to salt exposure. Our study establishes genomic and cellular resources for sand beans and provides a framework for investigating how natural variation in the regulation and cellular organization of conserved pathways contributes to environmental stress tolerance.

## Introduction

Soil salinization is an increasing threat to terrestrial ecosystems and agricultural productivity. The most recent global assessment by the Food and Agriculture Organization estimated that nearly 1.4 billion hectares, corresponding to slightly more than 10% of the global land area, are already affected by salinity, with an additional one billion hectares considered at risk because of climate change and unsustainable land and water management^1^. This problem is particularly urgent in coastal regions, where sea-level rise, storm surges, seawater intrusion, altered precipitation, and increasing freshwater demand can accelerate salinization of soils and groundwater^2–4^. Developing crops and crop relatives that can maintain growth in saline and coastal environments is therefore an important goal for climate-resilient agriculture.

Plant salt stress is typically characterized by osmotic stress, ion toxicity, nutrient imbalances, and secondary oxidative damage^5,6^. Plants respond through interconnected mechanisms, including osmotic adjustment, Na⁺ exclusion or compartmentation, K⁺ homeostasis, water transport, reactive oxygen species detoxification, hormone signaling, protein quality control, and growth regulation^5,7–9^. Much of the current mechanistic framework has been developed from Arabidopsis and major crops, where salt responses are often interpreted in terms of inducible pathways activated after stress perception. These include Ca²⁺-dependent signaling, the SOS pathway, HKT-mediated Na⁺ retrieval, ABA-associated responses, ROS signaling, and stress-responsive transcription factor networks^7,10–14^. This framework has been essential for understanding plant responses to salinity, but it may not fully explain how wild species tolerate salinity stress and adapt to coastal environments.

Naturally salt-adapted wild plants provide ideal systems to understand adaptive mechanisms that may be absent, reduced, or underused in crops^15–17^. Crop wild relatives and wild species often retain genetic and regulatory variation shaped by heterogeneous environments, making them valuable resources for improving abiotic stress tolerance^17–19^. Coastal populations are especially informative because they experience recurring combinations of salinity, salt spray, drought, high irradiance, nutrient limitation, and unstable sandy substrates^20–23^. Under such conditions, adaptation may depend not only on strong stress induction after exposure but also on constitutive or primed physiological states that provide protection before stress becomes severe^24,25^.

Sand bean, *Strophostyles helvola*, is a native North American wild legume species widely distributed across diverse habitats, including both coastal and inland areas (**Figure 1**). It is closely related to common bean (*Phaseolus vulgaris*^26^), a relationship we confirmed at the genome scale (**Figure 2**). Previous bulk-root transcriptomic work comparing salt-tolerant Beach and salt-sensitive Inland ecotypes, collected from coastal and inland habitats respectively, indicated that the two ecotypes differ substantially in their transcriptional responses to salinity^26^. However, bulk-tissue profiling averages expression across heterogeneous root cell populations, it cannot determine whether adaptive responses are localized to root-surface cells, diffusion barriers, vascular tissues, or proliferating cell states^27^. In addition, the lack of a chromosome-scale reference genome and cell-type-resolved transcriptomic resource has limited the use of sand bean as a system for studying the cellular and evolutionary basis of coastal adaptation.

**Figure 1.**
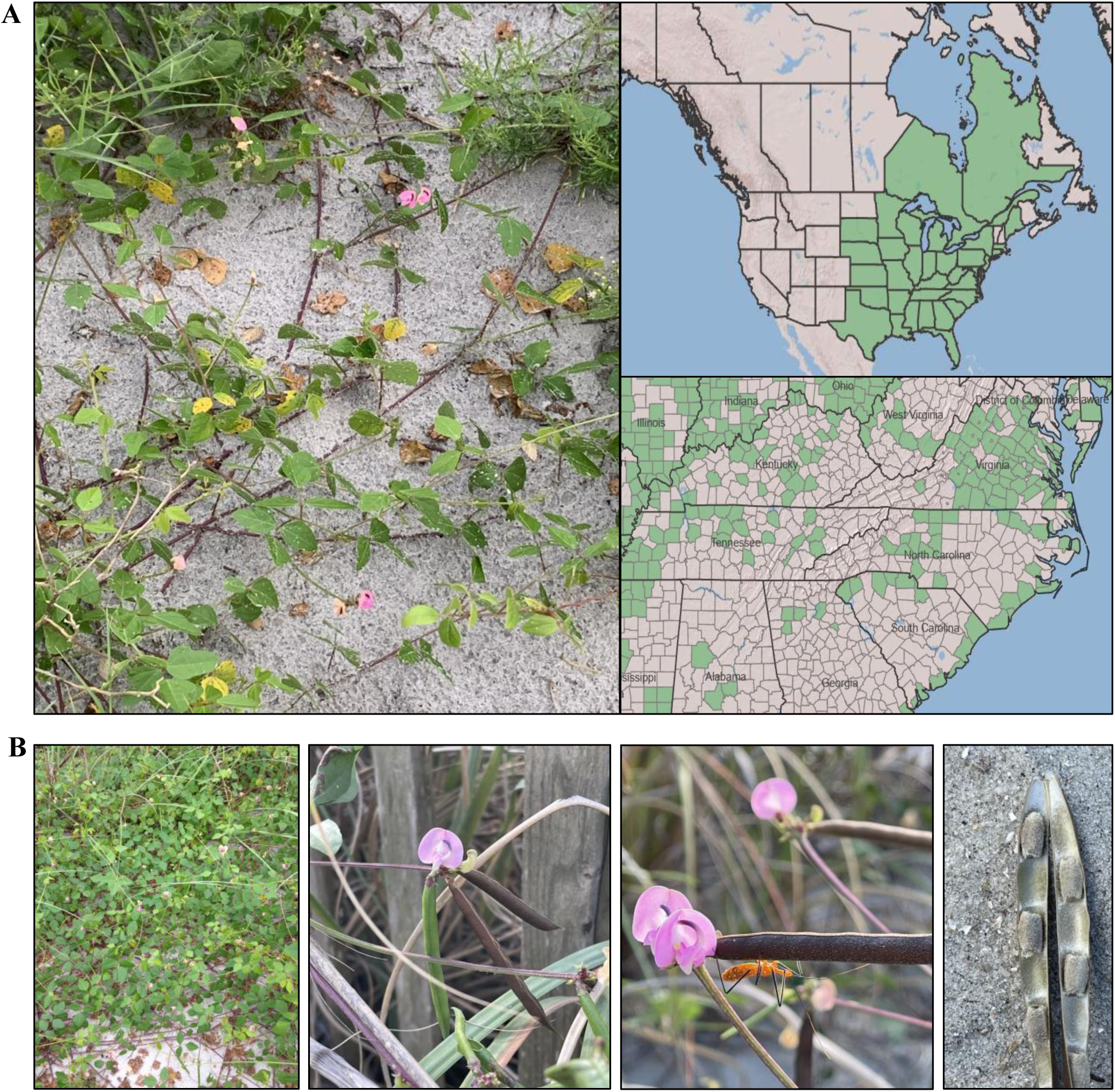
Geographic distribution, morphology and contrasting salt responses of Beach and Inland sand bean ecotypes. (A) Natural habitat and geographic distribution of sand bean *Strophostyles. helvola*. A representative plant is shown in a coastal sandy habitat. The maps show the distribution of sand beans in eastern North America and county-level occurrence records in the southeastern United States. Shaded regions indicate documented occurrences. (B) Representative morphological features of sand beans, including its prostrate-to-twining growth habit, trifoliate leaves, pink papilionaceous flowers, elongated pods and mature seeds.

**Figure 2.**
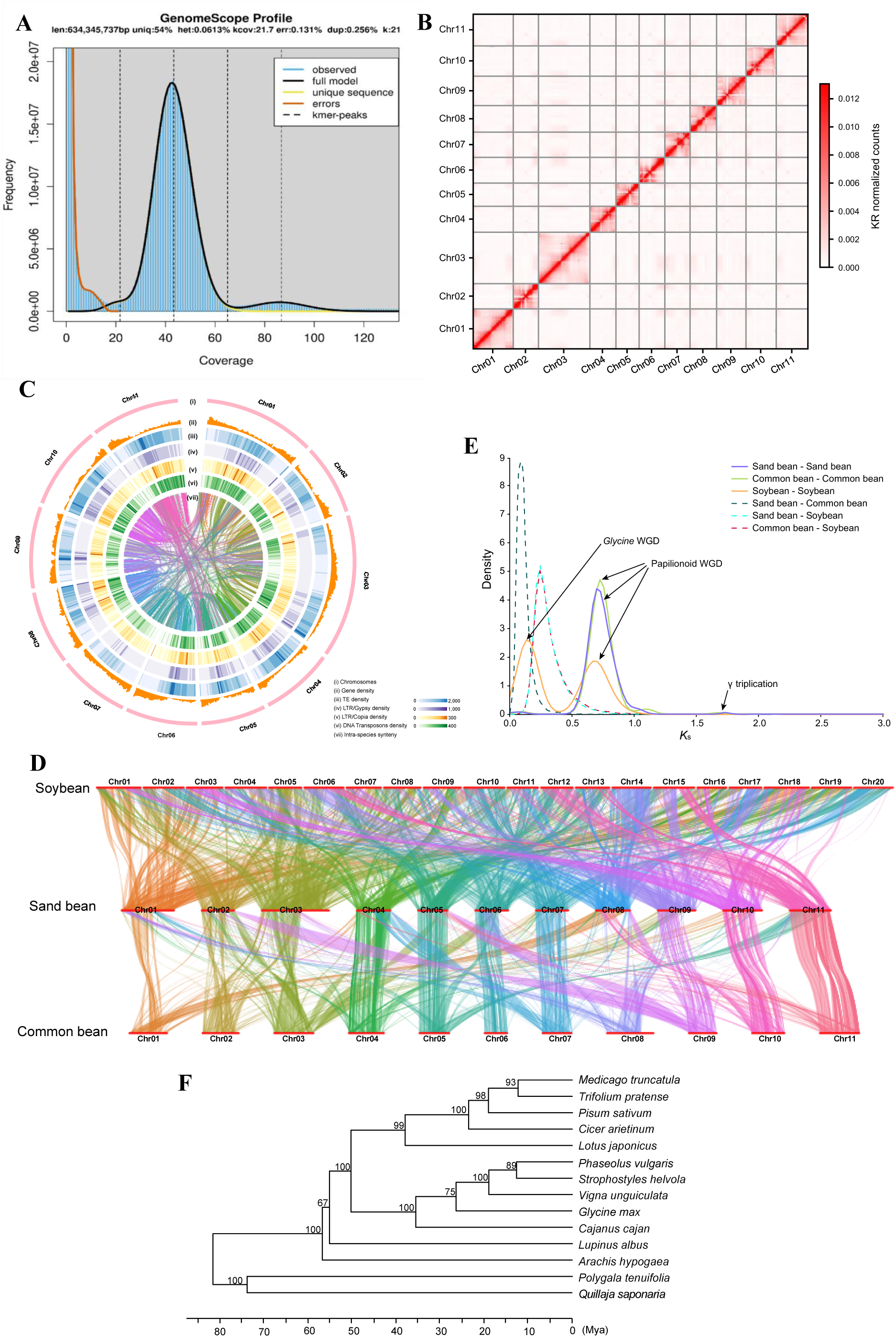
Chromosome-scale genome assembly and comparative genomic analyses of sand beans. **(A)** Estimation of sand bean genome size and heterozygosity based on k-mer frequency analysis. K-mer abundance was calculated from sequencing reads using Jellyfish with a k-mer size of 21, and genome size and heterozygosity were estimated using GenomeScope 2.0. **(B)** Hi-C contact map of the chromosome-scale *Strophostyles helvola* genome assembly. Interaction frequencies were calculated using 500-kb bins and visualized as a Knight–Ruiz-normalized contact matrix. Darker red indicates higher interaction frequency. **(C)** Genome landscape of *S. helvola*. Tracks show chromosomes, gene density, transposable element density, LTR/Gypsy retrotransposon density, LTR/Copia retrotransposon density, DNA transposon density, and intra-genomic syntenic relationships between duplicated genomic regions. Gene and repeat densities were calculated in non-overlapping 1-Mb windows. **(D)** Genome-wide syntenic relationships among soybeans (*Glycine max*), sand beans (*S. helvola*), and common beans (*Phaseolus vulgaris*). Syntenic blocks are represented by connecting lines, with colors corresponding to the 11 sand bean chromosomes. **(E)** Distributions of synonymous substitution rates (K_s) for paralogous and orthologous gene pairs within and among sand bean, common bean, and soybean. **(F)** Time-calibrated phylogenetic relationships among representative legume species. *Polygala tenuifolia* and *Quillaja saponaria* were used as outgroups. Estimated divergence times are indicated at the nodes, together with bootstrap support values.

Roots are composed of developmentally and functionally specialized cell populations. These cell types differ in their exposure to saline soil and in their roles in water uptake, ion transport, barrier formation, metabolism, growth, and long-distance signaling. Earlier cell-type-specific studies in Arabidopsis showed that root responses to high salinity depend strongly on cellular identity and developmental position^27^. Recent single-cell and single-nucleus transcriptomic studies have extended this principle to crop species roots^28–32^. These approaches make it possible to distinguish whole-root stress responses from those restricted to specific cell types. This distinction is particularly important for natural adaptation because evolutionary divergence may involve changes in where, when, and how strongly conserved pathways are deployed, rather than the evolution of entirely novel stress-response genes.

Despite rapid progress in single-cell genomics of plants, most root atlases have been generated for established model plants or major crops. Wild legumes adapted to coastal environments remain poorly represented in single-cell root resources. This gap limits our understanding of how natural selection reorganizes stress-response pathways at cellular resolution and hinders the identification of cell-type-specific adaptive programs relevant to crop salinity tolerance. Integrating a gene-annotated reference genome with a single-nucleus root atlas can connect ecological variation with transcriptional responses in naturally salt-adapted wild legumes and enable cellular-scale comparative framework for crop improvement.

Here, we generated a chromosome-scale, gene-annotated reference genome for *S. helvola* and used it to conduct comparative genomic analyses and construct a single-nucleus transcriptomic atlas of roots from the Beach and Inland ecotypes under control and salt-stress conditions. By combining the resulting evolutionary context with cell-type annotation, differential expression, functional enrichment, and comparisons of basal and salt-responsive expression, we sought to identify cell-type-specific transcriptional programs associated with natural variation in salt tolerance.

Specifically, we addressed four key questions: (i) how is the *S. helvola* genome organized, and what is its evolutionary relationship to closely related legume species; (ii) which major root cell types and cell states are present in sand bean, and how does salt stress alter their relative representation in the Beach and Inland ecotypes; (iii) which salt-responsive transcriptional programs are shared between the two ecotypes and which are ecotype- or cell-type-specific; and (iv) which root cell types and candidate regulatory modules show higher baseline activity of conserved stress-response programs in the Beach ecotype.

## Results

### Chromosome-scale genome assembly and comparative genomic analyses

To develop a chromosome-scale reference genome for sand bean, we generated 28.35 Gb of PacBio HiFi long-read sequences and 70.99 Gb of Hi-C data, representing approximately 44.7× and 111.9× coverage, respectively, of the sand bean genome, which has an estimated genome size of 634.3 Mb and a low heterozygosity of 0.061% (Fig. 2A). The final assembly was 602.2 Mb in length and comprised a total of 150 contigs with a contig N50 of 14.17 Mb. Using the Hi-C data, approximately 589.9 Mb (97.96%) of the assembled sequences were clustered into 11 pseudomolecules, corresponding to the 11 haploid chromosomes of sand bean. Hi-C contact maps displayed strong interaction signals concentrated along the main diagonal with clear chromosome boundaries, supporting accurate chromosome anchoring as well as contig ordering and orientation (Fig. 2B). The sand bean assembly achieved a BUSCO^33^ completeness score of 98.3% and an LTR Assembly Index (LAI)^34^ of 16.03. Further evaluation using k-mer analysis with Merqury^35^ revealed a consensus quality value (QV) of 57.6 and a k-mer completeness rate of 98.3% of the assembly. Collectively, these metrics demonstrate the high quality and completeness of the sand bean genome assembly.

A total of 27,731 protein-coding genes were annotated in the sand bean genome, of which 98.65%, 98.93%, and 84.33% had homologs in the GenBank nr, TrEMBL and Swiss-Prot databases, respectively. BUSCO analysis recovered 97.9% of conserved embryophyte genes in the annotated gene set, indicating a high level of annotation completeness. Repetitive sequences accounted for 55.47% of the assembled genome. Genome-wide analysis revealed that genes and transposable elements (TEs) were unevenly distributed across the chromosomes, with genes generally enriched toward both chromosome ends, whereas TEs were concentrated in the pericentromeric and central chromosomal regions (Fig. 2C). Intra-genomic synteny analysis revealed extensive duplicated chromosomal segments throughout the genome, consistent with the ancient whole-genome duplication (WGD) that occurred in the common ancestor of the Papilionoideae subfamily of legumes approximately 55 million years ago (Mya)^36^ (Fig. 2C).

To gain insights into the genome divergence and evolution, we first compared the sand bean genome with those of common bean (*Phaseolus vulgaris*) and soybean (*Glycine max*) within the Phaseolinae tribe. Gene collinearity analysis revealed extensive chromosome-scale synteny between sand bean and common bean, with most chromosomes showing a nearly one-to-one correspondence despite several large-scale structural rearrangements (Fig. 2D). In contrast, each sand bean chromosome corresponded to multiple soybean chromosomes, consistent with the lineage-specific WGD previously reported in soybean^37^. To further resolve the timing of genome duplication and species divergence, we analyzed synonymous substitution rate (*K*_s_) distributions for paralogous and orthologous gene pairs (Fig. 2E). The paralogous *Ks* distributions of sand bean and common bean showed a major peak at approximately *K*_s_ = 0.7, corresponding to the ancient Papilionoideae-specific WGD^36^. Soybean retained this ancestral WGD signal but also exhibited an additional peak at a lower *K*_s_ value, representing the more recent *Glycine*-specific WGD^37^. *Ks* distributions within and between these three species indicated that the ancient Papilionoideae WGD preceded the divergence of sand bean, common bean, and soybean, whereas the *Glycine*- specific WGD occurred after soybean diverged from the lineage leading to sand bean and common bean.

A time-calibrated phylogeny was reconstructed for representative legume species (Fig. 2F). The phylogeny recovered sand bean as the sister species of common bean within the sampled Phaseolinae, with an estimated divergence time of approximately 15 Mya, whereas the soybean lineage diverged earlier, approximately 20–25 Mya. These divergence estimates are consistent with the *K*_s_ distributions and chromosome collinearity analyses, collectively supporting a closer evolutionary relationship between sand bean and common bean than between sand bean and soybean.

### Construction of a single-nucleus transcriptome atlas of sand bean roots under salt stress

To investigate cell-type-resolved transcriptional responses associated with salt adaptation in sand bean (*Strophostyles helvola*), we generated single-nucleus RNA sequencing datasets from Beach and Inland ecotypes under control and salt-stress conditions. The experiment included five root datasets: Inland_Control (UNCC-1031-01); Beach_Control (UNCC-1031-02); Inland_SaltStressR1 (UNCC-1031-03); Inland_SaltStressR2 (UNCC-1031-04); Beach_SaltStress (UNCC-1031-05). Nuclei were isolated from pooled whole-root tissues, barcoded using the GEXSCOPE single-nucleus RNA-seq platform and sequenced using paired-end high-throughput sequencing by the SolusCell company (Fig. 3A). Across the five datasets, sequencing generated 1.74 billion reads (Supplementary Table S1). The percentage of reads with valid barcodes was highly consistent among samples, ranging from 86.50% to 86.93%. Genome mapping rates ranged from 84.62% to 85.12%, and 63.24%–63.48% of reads mapped to genes. After quality filtering, 87,901 high-quality nuclei were recovered, with 14,413–19,843 nuclei per sample. The median number of detected genes per nucleus ranged from 799 to 912, and total detected genes per sample ranged from 22,566 to 23,132. These results indicate that the five datasets were of comparable quality and suitable for integrated single-nucleus transcriptomic analysis.

**Figure 3.**
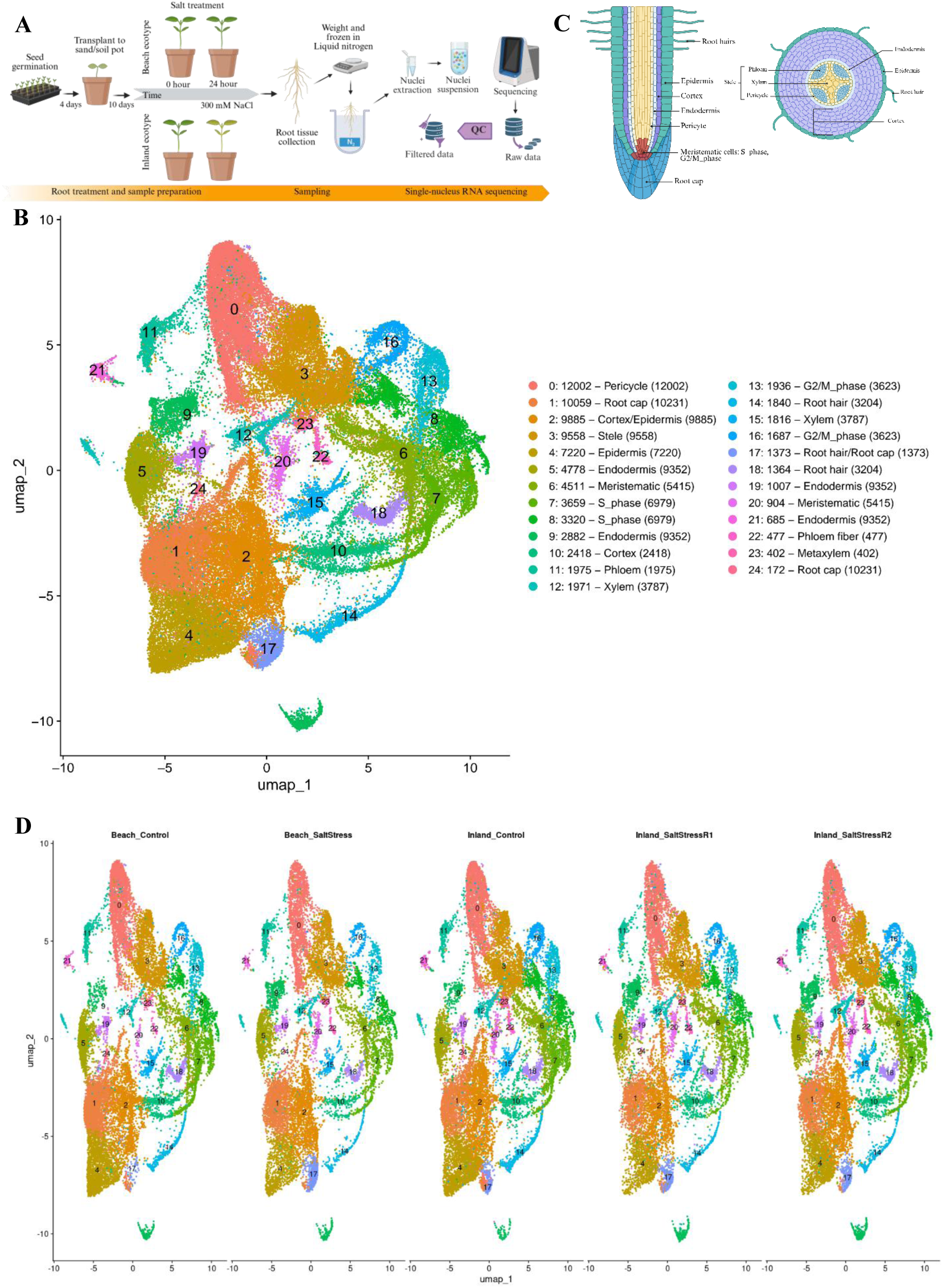
Construction and annotation of a single-nucleus transcriptomic atlas of sand bean roots. (A) Experimental design and single-nucleus RNA-sequencing workflow. Beach and Inland seedlings were maintained under control conditions or treated with 300 mM NaCl. Roots were collected 24 h after treatment and processed for nuclei isolation, library preparation, sequencing, quality control and generation of gene-expression count matrices by SolusCell. (B) UMAP representation of 87,901 high-quality root nuclei integrated across five datasets. Nuclei are colored by the 25 transcriptionally distinct clusters identified by graph-based clustering at a resolution of 0.4. (C) UMAP representation of the integrated atlas colored and labeled by the final cell-type annotations. (D) UMAP representations separated by dataset: Beach control, Beach salt, Inland control, Inland salt R1 and Inland salt R2. Nuclei are colored by their assigned cell types. R1 and R2 denote the two Inland salt datasets.

Integration of the five datasets resolved 25 transcriptionally distinct clusters, representing the cellular landscape of sand bean roots across both ecotypes under control and salt-stress conditions (Fig. 3B-C). Sample-split UMAPs showed that major cell populations were recovered across ecotypes and treatments, supporting the robustness of the integrated atlas (Fig. 3D). Cluster-level quality assessment further supported the robustness of the integrated atlas. Across most annotated clusters, UMI counts and the number of detected genes per nucleus showed broadly comparable distributions among five datasets (Fig. S1 and Fig. S2) indicating that major cell populations were not driven by strong sample-specific differences in sequencing depth or gene detection. Proliferating and meristematic clusters showed higher UMI and detected gene complexity than most differentiated cell types, consistent with their transcriptionally active cell states. These analyses established a single-nucleus transcriptomic atlas of sand bean roots, providing a foundation for subsequent cell-type annotation and functional characterization.

#### Identification and verification of root cell types

To annotate the 25 transcriptional clusters, we first identified cluster-specific marker genes (Supplementary Table S2 and Fig. S3), including the top 10 markers per cluster, which showed distinct cluster-restricted expression patterns (Fig. 3S). Because sand bean is a non-model legume with limited established root cell-type marker resources, we assigned cell type using Orthologous Marker Gene Groups (OMGs), a cross-species plant cell-type annotation framework that maps query sand bean marker genes to orthogroups and tests their overlap with cell-type markers of 15 reference species, to assign cell types^38^. OMGs returned a primary cell-type prediction and confidence score for each cluster (Fig. S4 and Supplementary Table S3), supported by orthogroup- overlap significance across the 15 reference species (Fig. S4).

OMG-based predictions were further validated using PlantScRNAdb^39^ by comparing sand bean cluster markers with homologous marker genes from soybean (*Glycine max*), *Medicago truncatula*, and *Arabidopsis thaliana* (Fig. S5; Fig. S6; Fig. S7; Supplementary Table S5). Soybean markers were prioritized as the closest available legume reference, whereas *Medicago* and *Arabidopsis* provided complementary evidence for cluster annotation. GO enrichment of cluster-enriched marker genes were consistent with the assigned cell identity for all clusters. Using this integrated framework, the 25 clusters were assigned to 16 major root cell types (Fig. 3B) (Supplementary note 1).

### Cell-type-specific responses to salt stress in Beach and Inland roots

Following cell-type annotation of the transcriptional clusters, we calculated the relative proportion of nuclei assigned to each annotated cell population within each sample (Figs. S8 and S9). Each sample contained multiple annotated populations, which were represented at unequal frequencies. Given the limited biological replication, these proportions are presented as descriptive summaries of the nuclei recovered from each sample.

We next identified salt-responsive differentially expressed genes (DEGs) within each annotated cell type (Fig. 4A and Supplementary Table S6). DEG numbers varied substantially among cell types, revealing pronounced heterogeneity in the transcriptional response. Downregulated DEGs generally outnumbered upregulated DEGs in both ecotypes, indicating extensive transcriptional repression and remodeling under salt stress. The largest responses in both ecotypes occurred in pericycle, endodermis, root cap, cortex/epidermis, stele, and epidermis (Fig. 4A). These populations also contained the largest sets of DEGs shared between Beach and Inland (Fig. 4B). These results identify barrier-associated, vascular-adjacent, root-tip, and outer-root tissues as major sites of conserved salt-responsive regulation.

**Figure 4.**
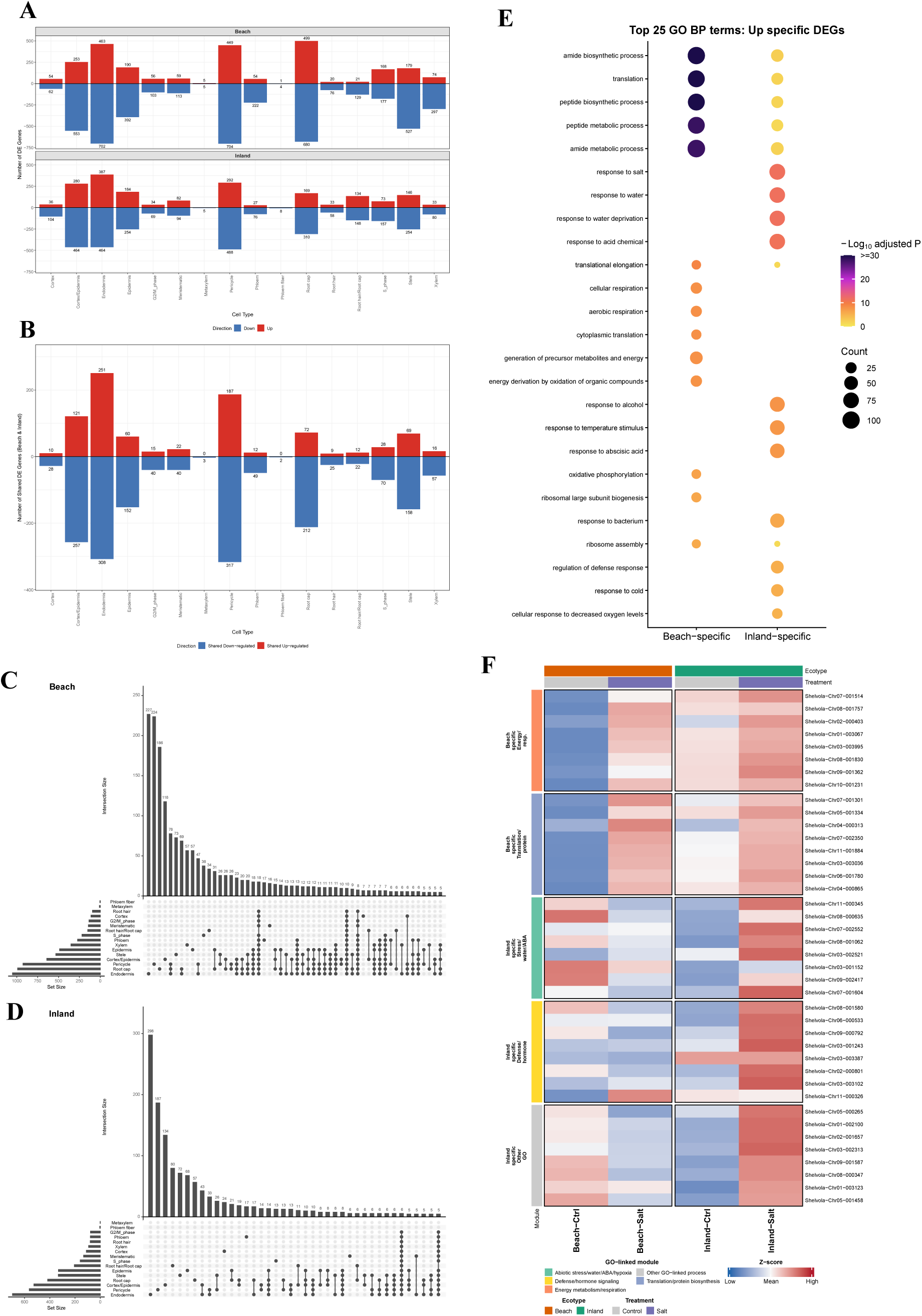
Cell-type-specific transcriptional responses to salt stress in Beach and Inland sand bean roots. (A-B) Numbers of salt-responsive differentially expressed genes (DEGs) across annotated root cell types in the Beach and Inland ecotypes. Salt-treated and control roots were compared separately within each ecotype and cell type. Upregulated and downregulated genes are shown as specific to one ecotype (A) or shared between the two ecotypes (B). (C) UpSet plot showing intersections among salt-responsive DEG sets across root cell types in the Beach ecotype. Horizontal bars show the total number of DEGs in each cell type, vertical bars show intersection sizes and the connected-dot matrix identifies the cell types included in each intersection. (D) UpSet plot showing intersections among salt-responsive DEG sets across root cell types in the Inland ecotype. Plot elements are defined as in B. (E) Gene Ontology (GO) enrichment analysis of ecotype-specific upregulated DEGs in Beach and Inland roots. Enriched Biological Process terms are shown on the y axis. Dot size indicates the number of genes assigned to each term, and dot color indicates the −log10-transformed adjusted P value. Terms with a Benjamini–Hochberg- adjusted P value < 0.05 were considered significant. (F) Heatmap showing the relative expression of representative ecotype-specific upregulated DEGs associated with the enriched Biological Process terms in D. Genes are grouped by their Beach- or Inland-specific classification and functional category. Columns represent Beach control, Beach salt, Inland control, Inland salt R1 and Inland salt R2; rows represent genes. Color indicates row-scaled average expression. For DEG analyses, significance was assessed using a two-sided Wilcoxon rank-sum test with Benjamini– Hochberg correction. Genes with an absolute average log2 fold change ≥ 0.25 and an adjusted P value < 0.05 were considered DEGs.

We then used UpSet analysis to determine whether salt-responsive DEGs were broadly shared across cell types or restricted to individual cell populations (Figs. 4C and 4D; Supplementary Table S7). In both ecotypes, the largest intersections consisted mainly of DEGs restricted to individual cell types. This result indicates that the root transcriptional response to salt stress is highly cell-type dependent. In Beach, the largest cell-type-specific DEG sets were detected in endodermis, root cap, and pericycle, followed by cortex/epidermis and stele. Thus, Beach deployed distinct transcriptional responses across barrier-associated, root-tip, root-surface, and vascular-adjacent populations. In Inland, the largest cell-type-specific set was detected in endodermis, followed by pericycle and cortex/epidermis. This distribution indicates a more concentrated, endodermis-centered response in Inland.

To define the functional differences between ecotypes, GO enrichment analysis of ecotype-specific upregulated DEGs identified both shared and contrasting functional programs (Fig. 4E and Supplementary Tables S8). Both ecotypes were enriched for translation (GO:0006412) and peptide biosynthetic process (GO:0043043), indicating a conserved protein-production response. Beach-specific genes were enriched for cellular respiration (GO:0045333), oxidative phosphorylation (GO:0006119), and generation of precursor metabolites and energy (GO:0006091). These results indicated greater induction of energy-producing processes in Beach than in Inland. In contrast, Inland-specific genes were enriched for response to salt (GO:1902074), response to water deprivation (GO:0009414), response to abscisic acid (GO:0009737), response to osmotic stress (GO:0006970), cellular response to hypoxia (GO:0071456), and response to oxidative stress (GO:0006979). Inland therefore showed stronger activation of canonical abiotic-stress pathways.

Cell-type-level GO analysis revealed how these functional programs were distributed across root populations (Fig. S10 and Supplementary Table S9). In Beach, the energy-metabolism signature was accompanied by enrichment of phenylpropanoid metabolism and plant-type secondary cell-wall biogenesis in individual cell types. This pattern suggests that energy production was coordinated with structural and metabolic remodeling. In Inland, salt-, water- deficit-, ROS-, and ethylene-related functions were especially represented in cortex/epidermis, epidermis, root hair, root cap, and endodermis populations. Thus, the Inland response was concentrated in outer-root, root-tip, and barrier-forming tissues that directly experience or regulate the effects of salt exposure.

Finally, a heatmap of representative DEGs underlying these enriched GO terms showed distinct ecotype-associated expression patterns across the four conditions (Fig. 4F). Genes selected from Beach-enriched categories generally showed higher relative expression after salt treatment in Beach roots. In contrast, genes selected from Inland-enriched categories showed stronger induction in salt-treated Inland roots. These gene-level patterns support a conserved translational response together with distinct ecotype-associated programs centered on energy and structural remodeling in Beach and canonical abiotic-stress signaling in Inland.

### Constitutive transcriptional programs associated with the Beach ecotype

To determine whether Beach roots possess transcriptional programs that are active before salt exposure and can be further reinforced under stress, we compared basal expression differences between the ecotypes with salt-responsive expression within Beach roots. Specifically, the log₂ fold change between Beach-Control and Inland-Control was compared with that between Beach- Salt and Beach-Control within each root cell type (Fig. 5A and Supplementary Tables S10 and S11). Quadrant I contained genes that showed both higher basal expression in Beach than in Inland and further induction following salt treatment in Beach. These genes represent pre-existing Beach- associated transcriptional states that can be reinforced after salt exposure. Quadrant IV genes also showed higher basal expression in Beach but were downregulated following salt treatment in Beach, representing Beach-associated baseline states that were attenuated rather than further activated under stress. The occurrence of both patterns across multiple root populations indicates that transcriptional preparedness in Beach roots is dynamically regulated in a cell-type-dependent manner.

**Figure 5.**
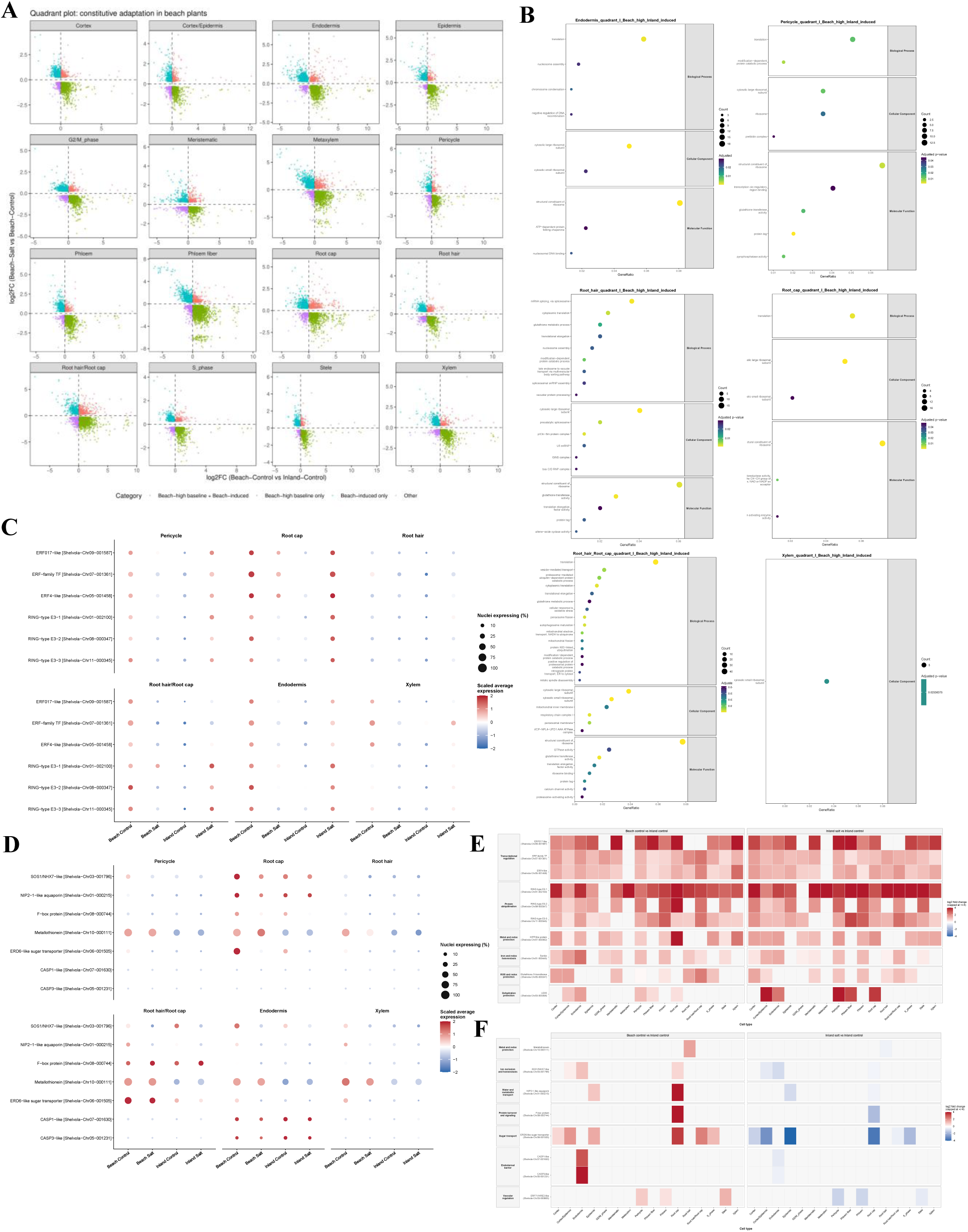
Cell-type-resolved constitutive and inducible expression patterns associated with Beach salt adaptation. (A) Quadrant plots relating basal expression differences between the ecotypes (x-axis, log₂ fold change of Beach-Control versus Inland-Control) to salt-responsive expression in Beach roots (y- axis, log₂ fold change of Beach-Salt versus Beach-Control) across 16 annotated root cell types. Quadrant I contains genes with higher basal expression in Beach that were further induced by salt in Beach. Quadrant II contains Beach-induced genes without higher basal expression in Beach. Quadrant III represents other expression patterns. Quadrant IV contains genes with higher basal expression in Beach that were downregulated after salt treatment in Beach. (B) Gene Ontology enrichment analysis of Quadrant I genes identified in (A). Significantly enriched biological processes are shown for six root cell populations. Dot position and size indicate gene count, and color represents −log₁₀(adjusted *P* value). (C-D) Expression of representative Quadrant I genes (C) and Quadrant IV genes (D) across four experimental conditions and six selected cell types. Dot size indicates the percentage of nuclei expressing each gene, and color represents scaled average expression. (E - F) Cell-type-resolved expression changes of representative Quadrant I (E) and Quadrant IV (F) genes across the 16 annotated root cell types. The left and right heatmaps show log₂ fold changes for Beach-Control versus Inland-Control and Inland-Salt versus Inland-Control, respectively. Genes are grouped by their predicted functions. Red and blue indicate positive and negative log₂ fold changes, respectively, with values capped at ±4.

To determine which biological functions were represented by genes that were already expressed at higher baseline levels in Beach roots and further induced by salt in Beach roots, we performed GO enrichment analysis for Quadrant I genes (Fig. 5B), focusing on six representative cell types, including endodermis, pericycle, root cap, root hair and xylem. Among these cell types, Root hair/Root cap displayed the broadest GO enrichment profile, suggesting that this root-surface transitional population is a major site where Beach baseline-enriched programs overlap with Inland salt-inducible responses. GO enrichment analysis showed that translation- and ribosome-associated terms were recurrent across multiple cell types, including endodermis, pericycle, root cap, root hair and Root hair/Root cap. Additional enriched terms were related to protein catabolism or ubiquitin-associated processes, glutathione metabolism, oxidative-stress response, vesicle and vacuolar transport, mitochondrial and peroxisomal functions, and chromatin- or nucleosome-associated processes. These functional categories suggest that genes already expressed at higher baseline levels in Beach roots include cellular maintenance, protein homeostasis, redox buffering and organelle-associated stress-response programs that are induced in Inland roots only after salt exposure.

Dot plots and heatmaps of representative genes across the four experimental conditions illustrated the contrasting expression patterns of the two groups (Fig. 5C-F). Several of these genes were concentrated in specific root populations, revealing a spatially organized component of the Beach-associated baseline program. At single-gene resolution, representative Quadrant I genes formed a recurrent regulatory backbone across multiple root cell types (Fig. 5C and Fig. 5E). The right-hand heatmap showed the Inland salt response (Inland salt versus Inland control), allowing the Beach baseline patterns to be compared directly with salt-induced expression in Inland (Fig. 5E-F). This group included the ERF-family transcription factors *Shelvola-Chr09-001587*, *Shelvola-Chr07-001361*, and *Shelvola-Chr05-001458*, together with three RING-type E3 ubiquitin ligases, *Shelvola-Chr01-002100*, *Shelvola-Chr08-000347*, and *Shelvola-Chr11-000345*, which were broadly detected across most root cell types. Additional Quadrant I candidates showed more restricted distributions, including the HIPP-like protein *Shelvola-Chr07-000362*, ferritin *Shelvola-Chr01-000445*, glutathione S-transferase *Shelvola-Chr05-000247*, and LEA5 *Shelvola- Chr03-003356*. Representative Quadrant IV genes exhibited stronger cell-type-associated distributions (Fig. 5D and Fig. 5F). In the root cap, representative genes included the SOS1/NHX7- like Na⁺/H⁺ exchanger *Shelvola-Chr03-001796*, the NIP2-1-like aquaporin *Shelvola-Chr01- 000215*, and the F-box protein *Shelvola-Chr08-000744*. Metallothionein *Shelvola-Chr10-000111* and the ERD6-like sugar transporter *Shelvola-Chr06-001505* were primarily detected in root-surface populations, whereas the CASP1- and CASP3-like genes *Shelvola-Chr07-001630* and *Shelvola-Chr05-001231* showed their strongest enrichment in the endodermis. The ERF71/HRE2- like transcription factor *Shelvola-Chr03-000805* was predominantly detected in vascular cell types.

Together, these analyses show that Beach roots combine elevated basal expression of stress- associated genes with their further reinforcement following salt exposure. This broadly distributed regulatory preparedness is accompanied by constitutive Beach-associated functions localized to specific root-surface, root-tip, endodermal, and vascular populations, revealing both shared and spatially organized components of coastal salt adaptation.

## Discussion

### Constitutive and cell-type-resolved transcriptional preparedness as a strategy for coastal salt adaptation

Plant salinity responses have traditionally been described as inducible processes initiated by salt perception and signal transduction, followed by transcriptional reprogramming and the activation of downstream protective responses^40,41^. Transcriptional profiling of salt-treated sand bean roots was broadly consistent with these conserved mechanisms. However, Beach and Inland roots differ substantially in the baseline regulation, cellular distribution and salt-induced reinforcement^41^. Compared with Inland plants, the Beach ecotype maintained higher baseline expression of numerous regulatory and protective genes while retaining the capacity for additional induction under salt stress. In our study, constitutive preparedness in Beach ecotype appeared to comprise at least two organizational layers: (i) elevated basal deployment of selected protective and regulatory programs while preserving their responsiveness to increasing salinity; and (ii) cell-type-specialized protective and transport modules.

The first layer integrated elevated baseline expression with preserved salt responsiveness. Beach roots showed elevated baseline expression of recurrent regulatory candidates, including ERF-family transcription factors and RING-type E3 ubiquitin ligases, while retaining the capacity for further salt-induced activation. The ERF017-like gene *Shelvola-Chr09-001587* was repeatedly detected among Beach-baseline-enriched candidates across multiple root cell types, whereas RING-type E3 ligase *Shelvola-Chr01-002100* recurred in cortex, endodermis, epidermis, meristematic, phloem, metaxylem and root-cap-associated populations (Fig. 5C and Fig. 5E). Their broad distribution suggests that ERF-mediated transcriptional regulation and ubiquitin-associated protein control may constitute shared regulatory components across distinct cell-type responses. Although the functionally characterized genes discussed below are not established one-to-one orthologs of the sand bean candidates, functional studies of other AP2/ERF transcription factors and RING-type E3 ligases support roles for these gene families in salt-stress regulation. In *Arabidopsis*, RAP2.6 binds stress-responsive cis-elements and regulates ABA-, salt- and osmotic- stress responses^42^, whereas soybean GmERF105 modifies the expression of LEA, ABA-associated and antioxidant genes under salinity^43^. Similarly, constitutive expression of the rice ERF transcription factor OsDRAP1 enhances salt tolerance and alters downstream genes involved in transcriptional regulation, ion transport and redox homeostasis^44^. RING-type E3 ligases similarly regulate several salt-response processes through selective protein ubiquitination. *Arabidopsis* STRF1 affects salt tolerance through membrane trafficking and ROS regulation^45^, whereas sweetpotato IbATL38 enhances salt tolerance when expressed in Arabidopsis and promotes stress-responsive transcription and ROS control^46^. Importantly, this elevated baseline state did not eliminate further salt responsiveness. Beach roots also retained the capacity to activate a subset of salt-responsive genes that were similarly induced in Inland roots upon salt exposure. Several Beach-baseline-enriched genes were additionally induced following salt exposure, including the LEA5 protein *Shelvola-Chr03-003356* and the heavy metal-associated isoprenylated protein *Shelvola-Chr07-000362*. These expression patterns suggest that dehydration protection and metal- or redox-associated functions were not only pre-positioned in Beach roots but could be further reinforced following salt exposure. Constitutive preparedness in the Beach ecotype therefore appears to represent a responsive pre-activated state rather than a fixed transcriptional endpoint. A similar expression strategy has been reported in the extremophyte *Thellungiella*, where several stress-associated genes, including *SOS1*, showed higher basal expression than in *Arabidopsis*^47^. *ThSOS1* remained responsive to salt, and reducing its expression strongly compromised salt tolerance, providing functional evidence that elevated basal activity of a key ion-homeostasis component can contribute to salt adaptation without eliminating subsequent stress regulation. Furthermore, comparative single-cell analysis of Brassicaceae roots revealed a closely related expression pattern in the extremophytes *Eutrema salsugineum* and *Schrenkiella parvula*^48^. A subset of stress-responsive orthologs showed high expression before salt exposure and remained highly expressed after treatment, at levels comparable to the salt-induced state of their stress-sensitive relatives. Wang et al. (2026)^48^ therefore described these genes as exhibiting a constitutively prepared expression state. This pattern parallels the elevated basal expression observed in Beach sand bean, although many Beach Quadrant I genes additionally showed further induction after salt exposure.

A second layer consisted of cell-type-specialized functional modules that were preferentially associated with distinct adaptive processes. Root cap and root hair cells may function as early protective interfaces with the saline environment. Beyond the general preparedness pattern, Wang et al. (2026) further showed that its deployment was not uniform across root cell types. Lateral root cap and atrichoblast populations contributed disproportionately to the prepared transcriptional state in the extremophytes, highlighting root-tip and outer-root cell populations as important sites of stress preparedness. This spatial organization provides a useful parallel to the Beach ecotype, in which elevated expression of selected protective functions was also concentrated in specific root cell populations. In our study, the Beach ecotype showed elevated expression of *SOS1/NHX7*-like Na^+^/H^+^ antiporter *Shelvola-Chr03-001796* together with the aquaporin *NIP2-1 Shelvola-Chr01-000215* in the root cap cells in the control condition, suggesting constitutive preparedness at root–soil interface (Fig. 5D and Fig. 5F). The importance of SOS1-mediated Na⁺ homeostasis is supported by previous findings in *Thellungiella*, where reduction of *ThSOS1* expression caused strong salt sensitivity and disrupted Na⁺ homeostasis^49^. Evidence from the coastal legume *Vigna marina* further links elevated basal *SOS1* expression with root Na^+^ excretion and salinity tolerance^50^. These findings support ion-homeostasis regulation at the root–soil interface as one component of the cell-type-specialized Beach state. Beach cortex and cortex/epidermis cells showed elevated expression of genes encoding a glutathione S-transferase (GST) *Shelvola-Chr05-000247*, ferritin *Shelvola-Chr01-000445*, LEA5 *Shelvola-Chr03-003356* and an ERD6-like sugar transporter *Shelvola-Chr06-001505* (Fig. 5F). The localization of an Arabidopsis ERD6 transporter to root cortex and epidermal cells^51^, together with previous detection of several GST isoforms during root epidermal development^52^, supports the plausibility of redox regulation and sugar transport in these outer root tissues. The elevated expression of ferritin and LEA5 further strengthened the potential contributions of iron homeostasis and dehydration protection to the Beach’s constitutive preparedness^53,54^. In the Beach endodermis, elevated expression of the CASP1- and CASP3-like genes (*Shelvola-Chr07-001630* and *Shelvola- Chr05-001231*) points to endodermal barrier regulation as another component of the cell-type-specialized response. Because CASP proteins are associated with Casparian strip formation, their expression pattern raises the hypothesis that Beach roots may adjust the endodermal barrier to regulate radial solute and ion movement toward the stele under salinity. Consistent with the importance of spatially restricted ion-transport regulation in inner-root tissues, studies in *Atriplex nummularia* support this spatial interpretation^55,56^: salt stress strongly increased plasma membrane H^+^-ATPase transcript accumulation in *Atriplex nummularia* roots, particularly in the epidermis of the root tip and the endodermis of the elongation and differentiation zones. This tissue-restricted response suggests that adaptive responses to salinity are preferentially deployed in root tissues that control ion uptake, radial transport, and entry into the vascular system. Consistent with this pattern, the ERF/HRE2-like gene *Shelvola-Chr03-000805* was preferentially associated with pericycle, phloem, and stele populations and showed a Beach-high Quadrant IV pattern in the stele, suggesting a distinct regulatory component of preparedness in the vascular compartment. Together, these patterns indicate that Beach-associated preparedness is spatially organized across the radial structure of the root, extending from the root–soil interface and endodermal barrier to inner vascular tissues.

In addition, constitutive preparedness in the Beach ecotype should not be equated with constitutive activation of the entire stress-response network. Maintaining a uniformly elevated stress state would be expected to impose substantial energetic and growth costs^57^. Studies in *Arabidopsis* have shown that improved salt tolerance can arise through attenuation of stress-associated growth inhibition rather than through stronger activation of all canonical stress pathways^9,14,58^. These studies demonstrated that successful stress acclimation requires coordination between cellular protection and the maintenance of growth-related processes. The Beach ecotype appeared to maintain selected protective and regulatory functions at higher baseline levels while retaining inducible capacity. Beach-specific salt-induced genes were enriched for respiration, oxidative phosphorylation, translation, and protein biosynthesis (Fig. 4E), suggesting that the response involved targeted metabolic and proteostatic reinforcement rather than broad activation of excessive stress responses. These functions may help sustain energy production, protein renewal, and cellular activity under salinity, thereby preserving resources required for developmental regulation. By contrast, Inland roots showed stronger enrichment for canonical responses to salinity, water deprivation, ABA, hypoxia, and oxidative stress (Fig. 4E), consistent with a larger reactive stress response. Thus, the greater magnitude of the Inland response should not be interpreted as evidence of superior tolerance. Instead, it may reflect the need to establish protective functions de novo after stress perception, potentially diverting cellular resources toward emergency stress signaling and away from ongoing metabolic and developmental processes. This interpretation is consistent with the dynamic framework described by Geng et al. (2013)^59^, in which salt adaptation involves coordinated regulation of stress responses and growth during successive phases of the response, rather than sustained activation of the entire stress-response network. Thus, the transcriptional differences observed between the Beach and Inland ecotypes reflect different regulatory strategies for deploying conserved salt-response programs, rather than simply differences in the magnitude of stress-responsive gene induction.

Thus, our findings do not replace the classical inducible-response model, but extend it by incorporating elevated basal deployment of selected protective modules and their cell-type-resolved organization into the framework of natural adaptation. Overall, Beach roots appear not to be maintained in a chronically stressed state, but instead to combine constitutive preparedness with spatially organized protective functions and targeted metabolic reinforcement, a strategy that may preserve developmental capacity and facilitate continued growth under salinity.

### A comparative working model for salt adaptation in Beach sand beans

Based on our findings, we proposed a working model in which Beach-associated salt tolerance is linked to a constitutively prepared and cell-type-specialized transcriptional state (Fig. 6). Beach and Inland roots use many of the same salt-response functions. However, these functions differ in their basal expression, distribution among root cell populations, and induction after salt exposure. Thus, the main difference between the two ecotypes may lie in the regulation and cellular organization of a shared stress-response system.

**Figure 6.**
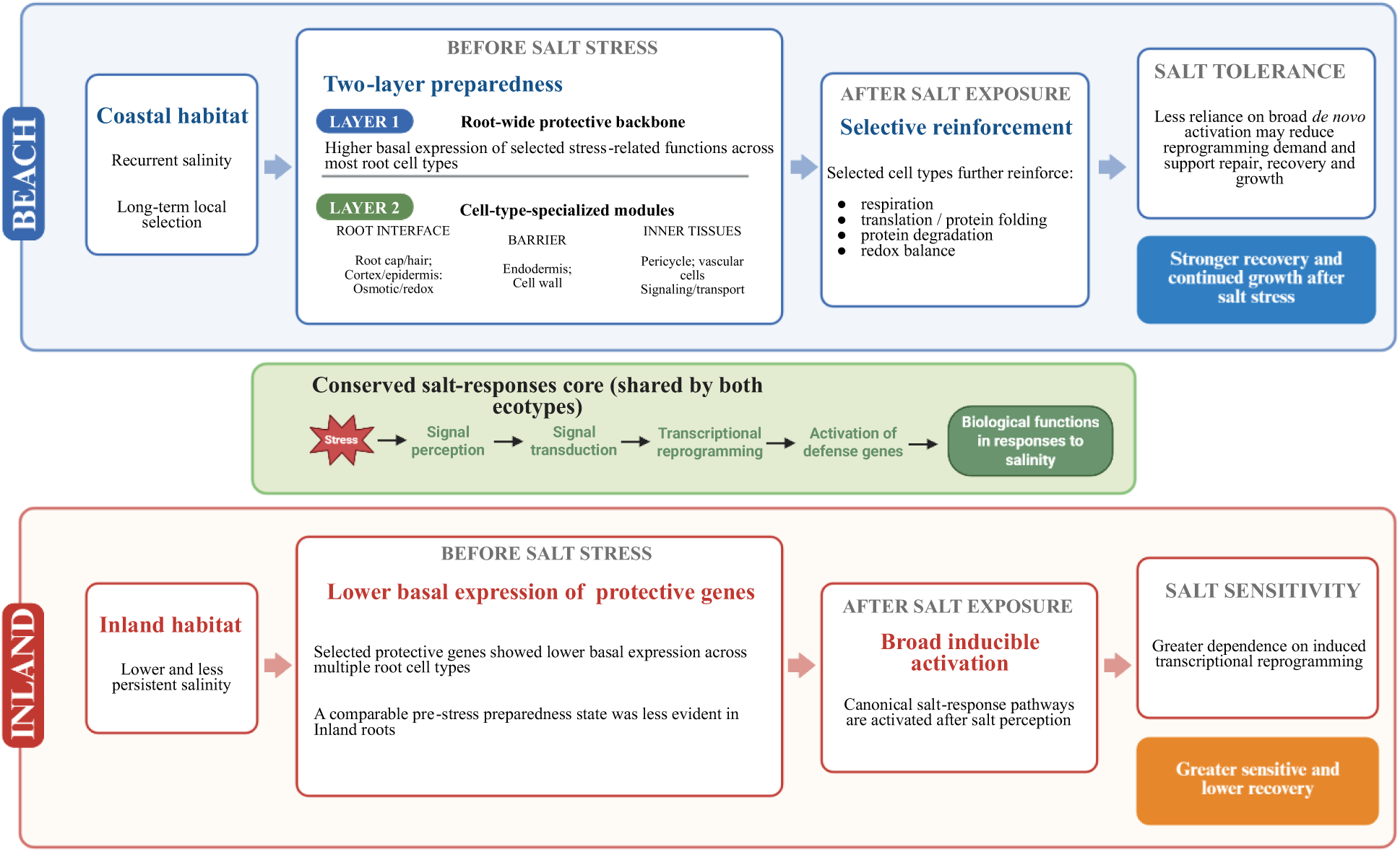
Working model for cell-type-resolved transcriptional preparedness associated with salt adaptation in the Beach ecotype. Beach and Inland sand beans share a conserved core salt-response system involving signal perception, signal transduction, transcriptional reprogramming and activation of protective functions. The model proposes that the ecotypes differ primarily in the basal deployment and subsequent induction of this shared response repertoire. Before salt exposure, Beach roots exhibit a two-layer preparedness state. The first layer consists of higher basal expression of selected stress- related functions across most root cell types. The second consists of cell-type-specialized modules distributed across complementary root compartments. Osmotic and redox functions are associated with cells at the root–soil interface, including root cap, root hair, cortex and epidermal cells. Cell- wall-related functions are positioned in the endodermal barrier, whereas signaling and transport functions are associated with the pericycle and vascular tissues. Following salt exposure, selected Beach cell populations further reinforce functions related to respiration, translation and protein folding, protein degradation and redox balance. Because part of the protective system is already active before stress, Beach roots may require less extensive *de novo* transcriptional activation, potentially reducing the demand for broad transcriptional reprogramming and supporting cellular repair, recovery and continued growth. This interpretation is consistent with the stronger post- stress recovery and growth observed in Beach plants. In Inland roots, selected protective genes show lower basal expression across multiple cell types, and a comparable pre-stress preparedness state is less evident. Inland plants may therefore rely more strongly on broad inducible activation of canonical salt-response pathways after stress perception, consistent with their greater salt sensitivity and lower recovery. Recurrent coastal salinity and long-term local selection are presented as an evolutionary hypothesis and were not directly tested. The proposed reduction in reprogramming demand and its contribution to recovery and growth also remain hypothetical. The model does not establish differences in response speed, energetic cost, or the causal contribution of any individual gene or cell population.

Plant species use diverse strategies to cope with salinity. For example, halophytes often possess specialized physiological or anatomical adaptations^60,61^. Different halophytes use combinations of strong ion exclusion, vacuolar ion sequestration, succulence, and external salt secretion through salt glands or epidermal bladder cells ^61,62^. These features reduce ion toxicity and help maintain cellular water balance during prolonged exposure to saline environments. In contrast, most crop species are glycophytes and often depend more strongly on reactive responses that are induced after salt exposure^5^. *Arabidopsis*-based frameworks commonly describe salt responses as progressing from stress perception and early signal transduction to transcriptional regulation and downstream protective processes, including ion homeostasis, osmotic adjustment, and redox regulation^40,41^. However, it places less emphasis on protective transcriptional states that are already present before stress occurs, which we propose here.

The Beach ecotype may use a complementary strategy. Rather than relying solely on specialized halophytic structures or a predominantly reactive transcriptional response, Beach may combine pre-existing transcriptional readiness with selective cell-type-specific reinforcement after salt exposure. In this model, selected genes associated with protective functions are expressed at higher baseline levels before stress, and a subset is further induced within specific root cell populations after exposure. This pattern may place Beach between specialized halophytes and salt- sensitive glycophytes that depend more strongly on inducible responses. Additional time points will be needed to determine the timing of this reinforcement.

This model is consistent with early comparisons between *Arabidopsis thaliana* and the extremophyte *T. halophila*^63^. Approximately 40% of their salinity-regulated transcripts were shared between the two species. However, the two species differed in the relative emphasis and regulation of these responses. *Arabidopsis* showed broad stress-induced transcriptional changes. In contrast, *Thellungiella* maintained a more stable cellular state and continued to grow under salinity. These findings showed that strong salt tolerance does not require completely new molecular pathways. It can also arise from different basal states and different regulations of conserved stress-response functions. Sand bean extends this concept to a within-species comparison at single-cell resolution. The Beach and Inland ecotypes are separated by a much narrower evolutionary distance than *Arabidopsis* and *Thellungiella*. Their contrasting tolerance may therefore reflect regulatory divergence within a largely shared genetic background.

Within-species ecological studies provide further support for this model. Coastal genotypes of *Panicum hallii* showed greater salt tolerance and better ion homeostasis than non-coastal genotypes^64^. They also showed tissue-dependent transcriptional responses to salinity. Later studies identified extensive constitutive and environmentally responsive expression differences among *P. hallii* accessions^65^. Many of these differences were associated with cis-regulatory variation. Coastal and inland ecotypes of *Mimulus guttatus* also differ in salt tolerance and fitness in their native environments^66^. These differences are accompanied by widespread cis- and trans-regulatory divergence and strong environmental effects on gene expression^67^. Together, these studies show that coastal adaptation can involve both stable expression differences and environmentally induced plasticity. They also indicate that regulatory divergence can contribute to local adaptation without requiring extensive changes in protein-coding sequences.

Recent cell-resolved studies provide a more direct cellular context for this model. In Brassicaceae extremophytes, prepared expression states were not distributed uniformly across root cell types. Different cell populations also contributed unequally to the salt response^48^. These results indicate that stress preparedness can emerge from the coordinated behavior of selected cell types. It does not require uniform activation across the whole root. A similar principle was found in wheat varieties with contrasting salt tolerance^30^. Root hair cells responded strongly in both varieties, but the varieties prioritized different cellular functions. The salt-sensitive variety showed stronger early signaling and osmotic adjustment, while the salt-tolerant variety placed greater emphasis on metabolic reprogramming and cellular repair. Sand bean showed a related pattern. In Beach roots, higher basal expression of genes associated with stress-related functions occurred across several annotated populations, including root cap and outer-root cells, endodermis, pericycle, and vascular-associated populations, rather than within one dominant population. These results suggest three connected features of Beach-associated preparedness: higher basal expression of selected genes, their nonuniform deployment among annotated cell types, and further induction of a subset after salt exposure. Inland roots retained many of the same inducible responses but showed lower basal expression of many genes that were already elevated in untreated Beach roots. We therefore propose that Beach combines pre-existing transcriptional readiness with targeted reinforcement in specific cell populations, whereas Inland depends more strongly on salt-induced transcriptional reprogramming.

### Sand bean as an emerging model for ecological and evolutionary systems biology

Beyond the biological insights into salt adaptation presented here, our study establishes sand bean as an emerging promising model for ecological and evolutionary systems biology in legumes. Several features make this species particularly attractive. First, sand bean is closely to common bean (*Phaseolus vulgaris*) (Fig. 2), findings in sand bean hold direct translational value for agriculture. By comparing naturally evolved traits and regulatory networks identified in sand bean with those of major legume crops, this framework offers a promising blueprint for discovering novel genetic mechanisms to enhance crop climate resilience. Second, sand bean has several biological and genomic features that favor its development as an experimental system. Its relatively compact genome is approximately half the size of the soybean genome, reducing the complexity of genome assembly, comparative genomics, and genetic analysis. Sand bean is also predominantly selfing, resulting in relatively high homozygosity within individuals and facilitating the development of genetically stable lines for genotype–phenotype studies. Its life cycle is manageable at approximately six months under current growth conditions and could potentially be shortened by optimizing controlled-environment growth and flowering conditions. Together, its compact genome, predominantly selfing mating system, and manageable generation time provide a practical foundation for functional genetic, genomic, and evolutionary studies. Third, sand bean can benefit from the extensive resources already developed for crop legumes. Its close evolutionary relationship with common bean, together with broader conservation across papilionoid legumes, enables comparative use of well-developed genome annotations, orthology relationships, molecular assays, and functional information from established legume systems. Candidate genes identified in sand bean can initially be evaluated through comparative and functional analyses of their orthologs in crops such as common bean and soybean, while species-specific genetic and molecular tools are developed. Thus, building a functional toolbox for sand bean does not need to begin from scratch. Our successful application of single-nucleus transcriptomics adds an important cellular dimension to this emerging system. The chromosome-scale reference genome and root cell atlas generated here provide a foundation for integrating population genomics, comparative genomics, single-cell transcriptomics, and other multi-omics approaches. Such integration will make it possible to move from genomic variation to cell-type-specific regulatory programs and, ultimately, to adaptive phenotypes.

Moreover, importantly, sand bean offers an ecological dimension that conventional crop models cannot easily provide. It has a broad geographic distribution across eastern and central North America and occupies strikingly diverse habitats, ranging from saline coastal dunes to inland Piedmont and mountainous environments. These naturally occurring populations provide replicated evolutionary experiments in which different environmental pressures may have shaped distinct adaptive strategies. Sampling populations across these environmental gradients will enable direct investigation of how genomic variation, regulatory programs, and cell-type-specific responses are associated with local environments.

### Future perspectives

Future studies should extend the present comparison of two representative ecotypes to broader coastal and inland populations to determine how consistently the transcriptional patterns identified here are associated with environmental variation. Integrating population genomics and pangenomics with cell-type-resolved transcriptomics will help identify adaptive alleles, cis-regulatory variation, and structural variants underlying ecotype-associated expression patterns. Functional validation using gene editing, transgenic approaches and spatial expression analyses of candidate genes, together with physiological measurements of ion homeostasis, barrier function and water relations, will be needed to establish causal links between candidate regulators, cell- type-specific programs, and salt tolerance. Because coastal habitats impose a complex combination of environmental challenges, including salinity, drought, nutrient limitation, and wind exposure. Future studies should therefore determine whether the transcriptional preparedness observed in the beach ecotype is primarily associated with salinity or represents a broader adaptive strategy for life in coastal environments. Together, these directions will connect natural environmental variation with genomic evolution, cellular regulation, and adaptive phenotype.

In conclusion, this study provides a chromosome-scale genome and single-nucleus root atlas for sand beans and establishes a cell-type-resolved framework for understanding coastal salt adaptation. Rather than identifying an entirely distinct salt-response toolkit, our results suggest that ecological divergence is associated with differences in the regulation and cellular organization of a largely conserved stress-response programs. This framework establishes *Strophostyles helvola* as a promising system for investigating the genomic and cellular basis of environmental adaptation in wild legumes. More broadly, this framework provides a foundation for determining how naturally evolved regulatory strategies can inform the discovery and eventual translation of adaptive mechanisms for improving stress resilience in legume crops.

## Methods and Materials

### Plant materials and salt treatment

Beach and Inland ecotypes of sand bean (*Strophostyles helvola*) were used in this study. The Beach ecotype represents a salt-tolerant coastal population, whereas the Inland ecotype represents a salt-sensitive inland population. Seeds were surface-disinfected with 5% sodium hypochlorite for 5 min, scarified, and germinated in sterilized vermiculite under controlled growth-chamber conditions at 27°C with a 16-h light photoperiod. After 4 days of germination, seedlings were transplanted into 4-inch clay pots containing sterilized 1:1 sand/soil mixture. Plants were watered daily, except on the treatment day and the tissue collection day. For salt treatment, 10 days post-transplantation, seedlings were treated with salt solution by applying 30 mL of 300 mM NaCl solution per plant at 10:30 AM. Control plants were treated with 30 mL of water. The following day at 10:30 AM, 24 h after treatment, root tissues were collected, flash-frozen with liquid nitrogen, and stored at −80 °C until needed.

Five root datasets were generated for single-nucleus RNA-seq: Inland_Control (UNCC- 1031-01), Beach_Control (UNCC-1031-02), Inland_SaltStressR1 (UNCC-1031-03), Inland_SaltStressR2 (UNCC-1031-04), and Beach_SaltStress (UNCC-1031-05). Inland_SaltStressR1 and Inland_SaltStressR2 are biological replicates. Each sample consisted of three pooled replicates, with approximately 500–600 mg total root tissue per sample. Roots were carefully removed from pots, rapidly rinsed with deionized water to remove soil particles, blotted dry with paper towels, weighed, immediately flash frozen in liquid nitrogen, and stored at −80°C until nuclei extraction for snRNA-seq.

### Genome assembly, annotation and comparative genomic analyses

PacBio HiFi reads were first processed using HiFiAdapterFilt^68^ (v2.0.1) to remove residual adapter sequences and then assembled into contigs using hifiasm^69^ (v0.14.2-r315). Hi-C reads were processed using Trimmomatic^70^ (v0.39) to remove adapter contamination and low-quality sequences and were subsequently aligned to the assembled contigs using BWA-mem^71^ (v0.7.17) with default parameters. Chromosome-scale pseudomolecules were generated using Juicer^72^ and 3D-DNA^73^, followed by manual curation. Hi-C contact maps were generated using HiCExplorer^74^ (v3.0) to assess chromosome anchoring and contig ordering and orientation. Gene space completeness was assessed using BUSCO^33^ (v5.3.2) with the embryophyta_odb10 database. Base-level accuracy and assembly completeness were evaluated using Merqury^35^ (v1.3) with the HiFi reads. Assembly quality was further assessed using the LTR Assembly Index (LAI)^34^ (v2.9.7).

Repetitive sequences in the sand bean genome were identified using EDTA^75^ (v2.0.1). Protein-coding genes were predicted using Helixer^76^ (v0.3.2), a deep learning-based gene prediction framework that integrates neural networks with hidden Markov models. The completeness of the predicted gene set was assessed using BUSCO^33^ (v5.3.2) with the embryophyta_odb10 database. Functional annotation of the predicted genes was performed by searching their protein sequences against GenBank nr and UniProt (TrEMBL and Swiss-Prot) databases using diamond^77^ (v2.1.11) with an e value cutoff of 1e-5.

Genome-wide syntenic blocks between sand bean and common bean, and between sand bean and soybean, were identified using MCScanX^78^ (v1.0.0) with default parameters. Synonymous substitution rates (*K*_s_) of paralogous and orthologous gene pairs were calculated using quota_Anchor^79^ (v1.0.2). For phylogenetic analysis, single-copy orthologous genes identified by OrthoFinder^80^ (v2.5.5) from representative legume species were used to reconstruct the species tree. Divergence times were estimated using BEAST2^81^ under an uncorrelated log-normal relaxed molecular clock with a Yule tree prior. Two calibration constraints were applied: the divergence between Fabaceae and *Quillaja saponaria* (75–85 Mya)^82^ and the divergence between *Glycine max* and *Lotus japonicus* (45–50 Mya)^83^.

### Nuclei extraction, single-nucleus RNA-seq library preparation and sequencing

Root samples were processed by SolusCell using a proprietary nuclei extraction method. Nuclei quality was assessed by microscopy-based examination of nuclear morphology and RNA-profile evaluation to ensure suitability for single-nucleus RNA-seq library preparation.

Single-nucleus RNA-seq libraries were prepared using Singleron Biotechnologies’ GEXSCOPE® platform, which includes nuclei capture, barcoding and library construction. Libraries were sequenced using paired-end, high-throughput sequencing on Illumina platforms.

### Single-nucleus RNA-seq preprocessing and generation of count matrices

Raw FASTQ files were processed using SolusCell’s internal single-nucleus RNA-seq workflow, which is based on CeleScope (https://github.com/singleron-RD/CeleScope) from Singleron Biotechnologies. Reads were aligned to the annotated *Strophostyles helvola* reference genome, the genome and annotation are available at http://bioinfo.bti.cornell.edu/lab/S_helvola.tar, with a total of 27,731 genes predicted achieving a BUSCO completeness score of 97.9%. Both raw and filtered count matrices were generated and provided for each sample. The raw matrix represents the initial gene-by-cell expression output from the single-nucleus RNA-seq workflow, whereas the filtered matrix contains high-quality nuclei retained after quality-control filtering. Raw and filtered matrices were shared with the laboratory through Globus. For all downstream analyses in this study, filtered matrices were used as input. The five single-nucleus RNA-seq datasets used in this study were organized as follows: Inland_Control (UNCC-1031-01); Beach_Control (UNCC-1031- 02); Inland_SaltStressR1 (UNCC-1031-03); Inland_SaltStressR2 (UNCC-1031-04); Beach_SaltStress (UNCC-1031-05)

### Quality control, integration, clustering and visualization

Likely doublets are identified with scDblFinder^84^ (v1.10.0) with default parameters and removed before downstream analysis^84^. After quality filtering, the five-dataset atlas comprised 87,901 high-quality nuclei, with 14,413–19,843 nuclei recovered per sample. The median number of detected genes per nucleus ranged from 799 to 912 across samples, and total detected genes per sample ranged from 22,566 to 23,132. Filtered gene-by-cell matrices generated after were then imported into R 4.5.1 and analyzed using Seurat v5.0.0^85^.

To normalize gene expression across nuclei, we applied the “NormalizeData” function using the LogNormalize method with a scale factor of 10,000^86^. Highly variable genes were identified using the “FindVariableFeatures” function, and the top 2,000 variable genes were retained. Then, to integrate the five datasets and reduce sample-specific effects, we used reciprocal principal component analysis-based integration in Seurat. Integration features were selected using the “SelectIntegrationFeatures” function, and each sample was scaled and subjected to principal component analysis before anchor identification. Integration anchors were identified using the “FindIntegrationAnchors” function with reciprocal PCA reduction, and the integrated expression matrix was generated using the “IntegrateData” function^87^. To integrate the five datasets and reduce sample-specific effects, we used reciprocal principal component analysis-based integration in Seurat. Integration anchors were identified using the “FindIntegrationAnchors” function with reciprocal PCA reduction, and the integrated expression matrix was generated using the “IntegrateData” function. Graph-based clustering was performed using the “FindClusters” function with a resolution of 0.4 and the Louvain algorithm^88^. This analysis resolved 25 transcriptionally distinct clusters. UMAP visualization was performed using the “RunUMAP” function based on the first 45 reciprocal PCA dimensions, with min.dist = 0.3, n.neighbors = 30 and cosine distance^89^. UMAP plots were generated to display sample identity, Seurat cluster identity and final cell-type annotations.

For each cluster, sample contribution was summarized as both absolute nuclei number and relative percentage. Sample-split UMAPs and composition summaries were used to evaluate cluster distribution across datasets and to compare cell-type composition between Inland salt stress replicate 1 and Inland salt stress replicate 2. Filtered matrices from all five datasets were used to construct the sand bean root cell atlas.

Cluster-enriched marker genes were identified using the “FindAllMarkers” function^90^. Only positively enriched marker genes were retained, and genes were required to be expressed in at least 10% of nuclei in the target cluster with |log_2_ fold change| ≥ 0.25. Marker genes with adjusted *p_value* < 0.05 were considered significant for downstream interpretation. In addition, the top 30 markers per cell type were selected for downstream visualization and annotation support (Supplementary Table S2).

### Cell-type annotation

Cell-type clusters were annotated using the Orthologous Marker Gene Groups (OMGs) pipeline^38^ (Sandbean branch, https://github.com/LiLabAtVT/OrthoMarkerGeneGroups), which includes Strophostyles helvola in its OrthoFinder-derived orthogroup table, allowing direct ortholog mapping of sand bean genes. Cluster-enriched marker genes from FindAllMarkers were used as input, and the top 200 genes per cluster, ranked by average log₂ fold change, were mapped to orthogroups and tested for overlap with reference cell-type marker groups using a one-sided Fisher’s exact test with Benjamini–Hochberg correction at an FDR threshold of 0.01. Final cell-type labels were assigned from the consolidated OMG prediction for each cluster, together with cluster-enriched marker-gene expression patterns. Orthogroup assignments across 35 species used in the OMG analysis are provided in Supplementary Table S4.

### Identification and enrichment analyses of DEGs

Differentially expressed genes (DEGs) were identified separately within each annotated cell type by comparing salt-treated and control samples in the Beach and Inland ecotypes. Genes with an absolute log2 fold change ≥ 0.25 and an adjusted *P* value < 0.05 were considered significant DEGs. Upregulated and downregulated genes were classified according to the direction of the log2 fold change^86^. Gene Ontology enrichment analysis was conducted separately for selected DEG and quadrant-derived gene sets using clusterProfiler package in R^91^. GO terms with adjusted *P* values < 0.05 were considered significantly enriched and visualized using dot plots.

## Acknowledgments

We gratefully acknowledge funding support from the University of North Carolina at Charlotte for this project. The Song Lab also acknowledges general research support from the National Science Foundation (grant no. 2318746), the National Institutes of Health (grant no. 1R15AT011603-01A1), and the North Carolina Biotechnology Center (grant no. FLG- 3806).

## Author Contributions

Conceptualization: BHS; Methodology: BHS, ZF, MK, PY, QX, SL; Experiments: PY, QTL; Analysis: XZ, QTL, HWS, TC; Field and Images: QX and BHS; Writing – original draft: QTL and BHS; Writing – review & improvement: All authors; Supervision: BHS and ZF.

## Conflicts of Interest

The authors declare no conflict of interest.

## Notes

### Competing Interest Statement

The authors have declared no competing interest.

