## Supplemental Figures for "A chromosome-scale reference genome and single-nucleus root atlas reveal the cellular and molecular basis of coastal adaptation in sand bean (*Strophostyles helvola*)"

### Supplementary data

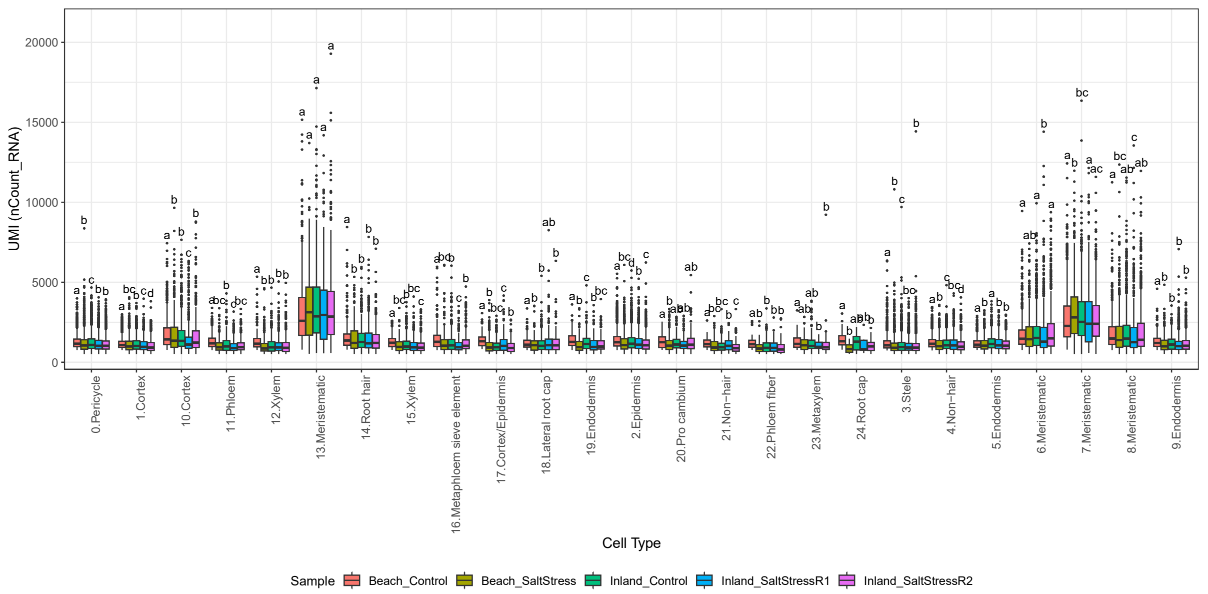


**Figure S1. Distribution of UMI counts across sand bean root clusters and datasets.**

Boxplots show the total number of unique molecular identifiers detected per nucleus (nCount_RNA) for each cluster–cell-type combination across Beach-Control, Beach-SaltStress, Inland-Control, Inland-SaltStressR1, and Inland-SaltStressR2. Colors indicate individual datasets. Lowercase letters above the boxplots denote multiple-comparison groupings among datasets within each cluster.

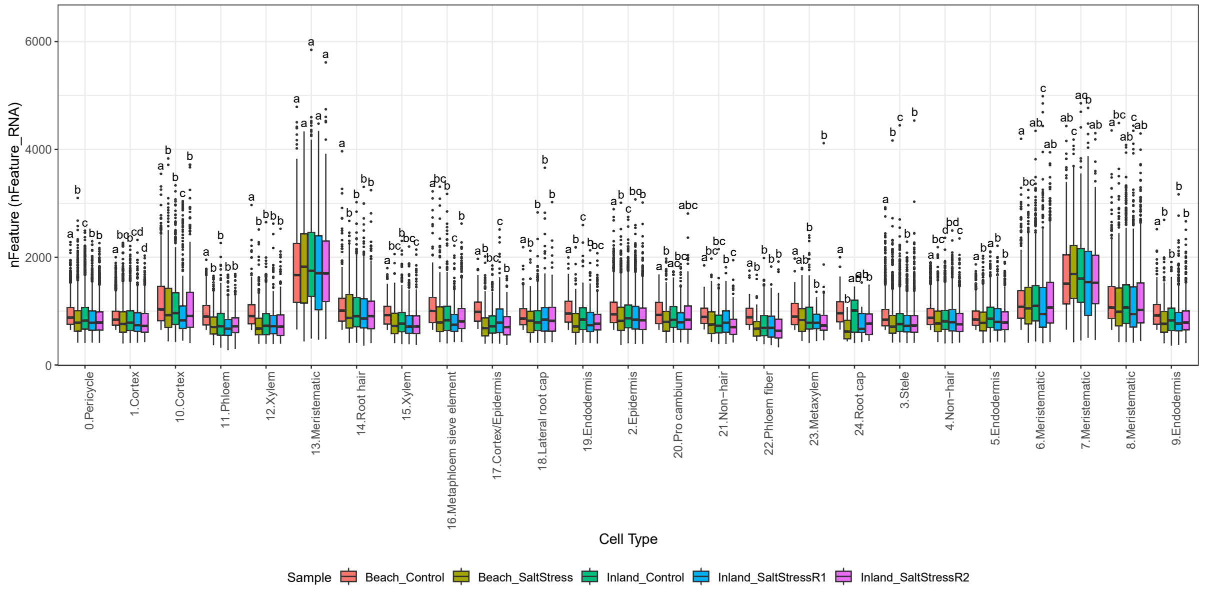


**Figure S2. Distribution of detected genes across sand bean root clusters and datasets.**

Boxplots show the number of genes detected per nucleus (nFeature_RNA) for each cluster–cell-type combination across the five datasets. Colors indicate Beach-Control, Beach-SaltStress, Inland-Control, Inland-SaltStressR1, and Inland-SaltStressR2. Lowercase letters denote multiple-comparison groupings among datasets within each cluster.

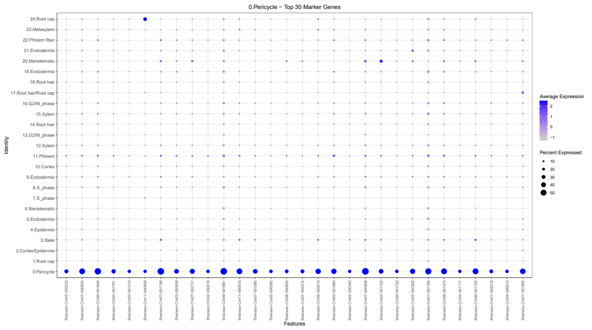
0.Pericycle - Top 30 Marker Genes
Identity
Features
Average Expression
Percent Expressed

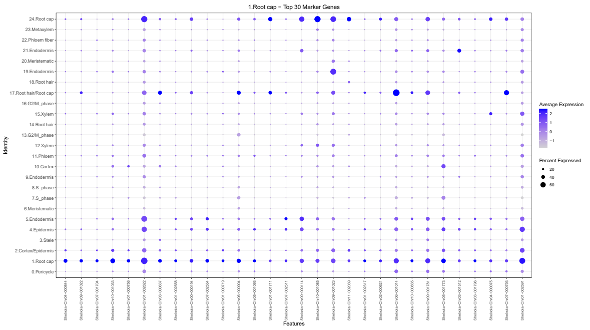
1.Root cap - Top 30 Marker Genes
Identity
Features
Average Expression
Percent Expressed

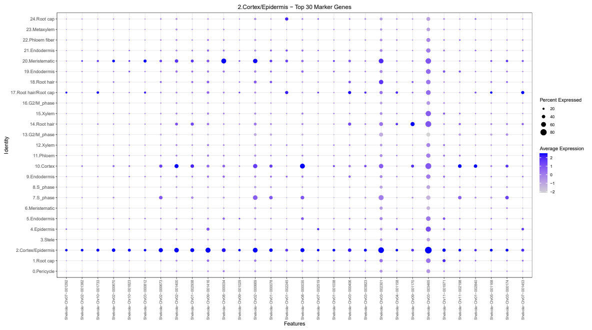
2.Cortex/Epidermis - Top 30 Marker Genes
Identity
Features
Percent Expressed
Average Expression

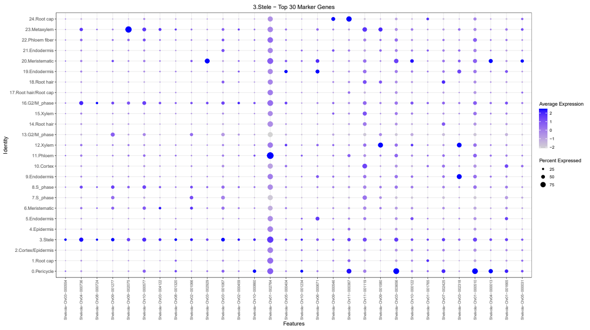
3.Stele - Top 30 Marker Genes
Identity
Features
Average Expression
Percent Expressed

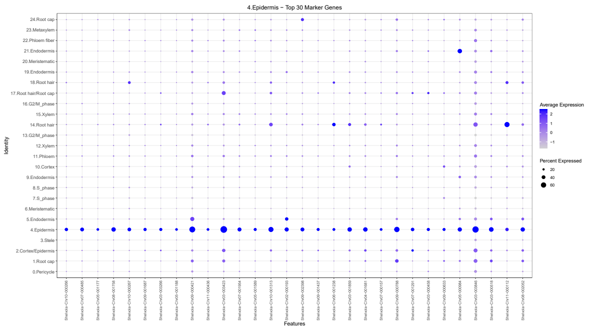
4.Epidermis - Top 30 Marker Genes
Identity
Features
Average Expression
Percent Expressed

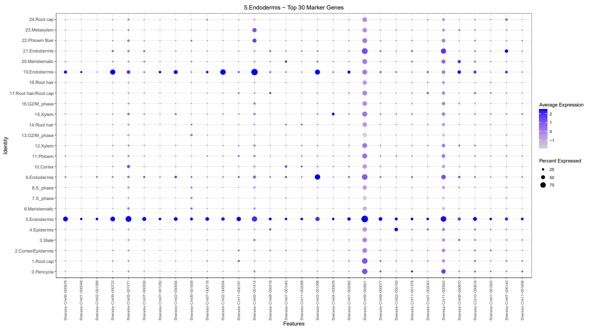
5.Endodermis - Top 30 Marker Genes
Identity
Features
Average Expression
Percent Expressed

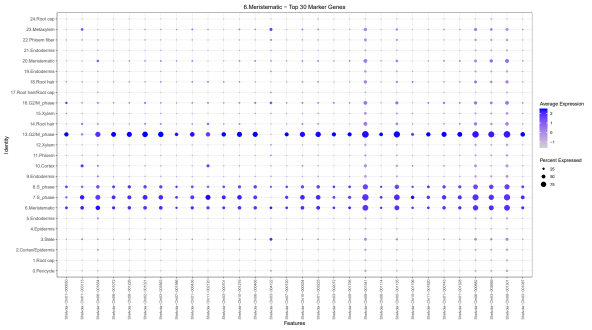
6.Meristematic - Top 30 Marker Genes
Identity
Features
Average Expression
Percent Expressed

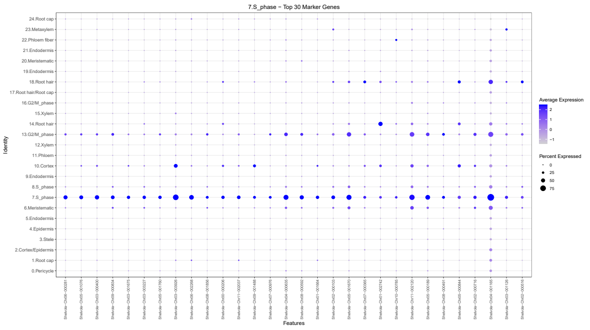
7.S_phase - Top 30 Marker Genes
Identity
Features
Average Expression
Percent Expressed

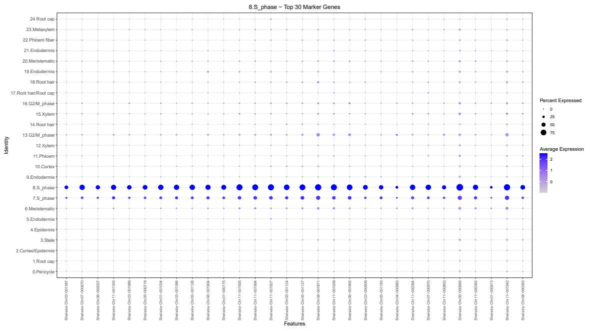
8.S_phase - Top 30 Marker Genes
Identity
Features
Percent Expressed
Average Expression

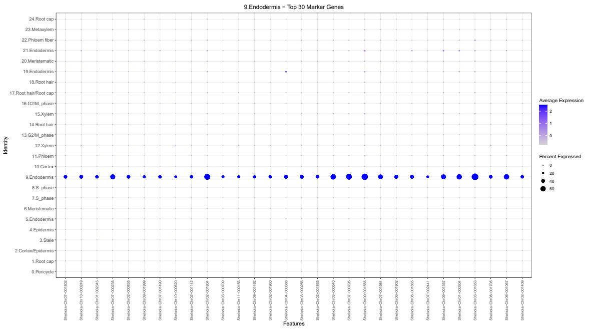
9.Endodermis - Top 30 Marker Genes
Identity
Features
Average Expression
Percent Expressed

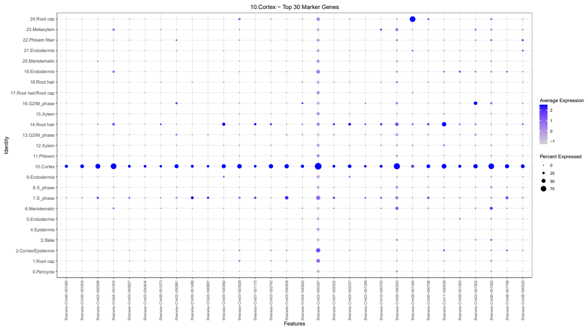
10.Cortex - Top 30 Marker Genes

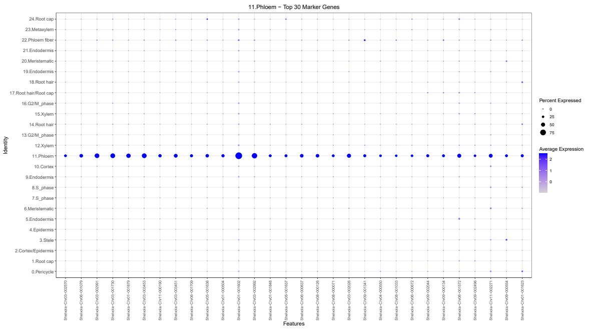
11.Phloem - Top 30 Marker Genes

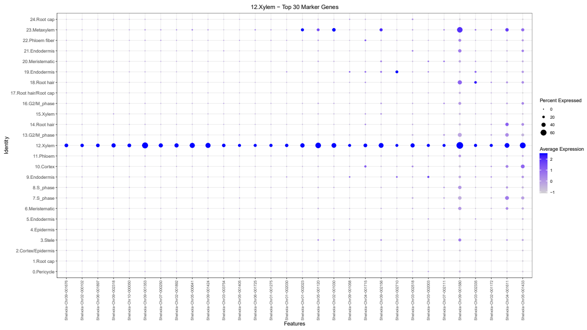
12.Xylem - Top 30 Marker Genes

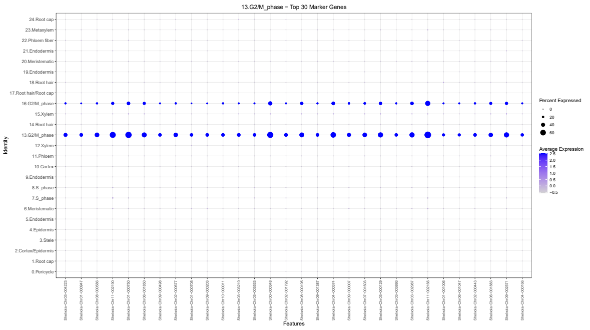
13.G2/M_phase - Top 30 Marker Genes

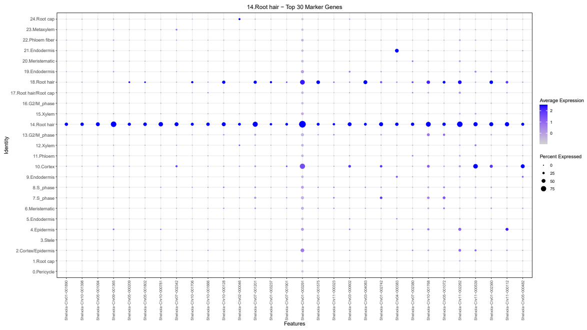
14.Root hair - Top 30 Marker Genes

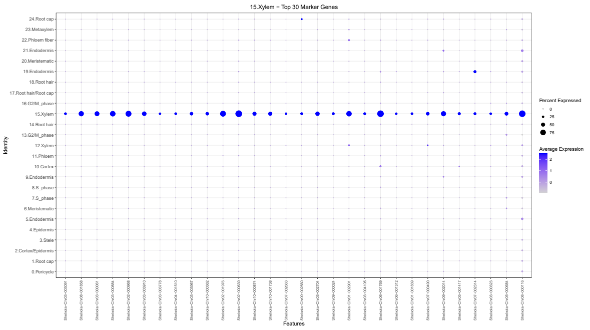
15.Xylem - Top 30 Marker Genes

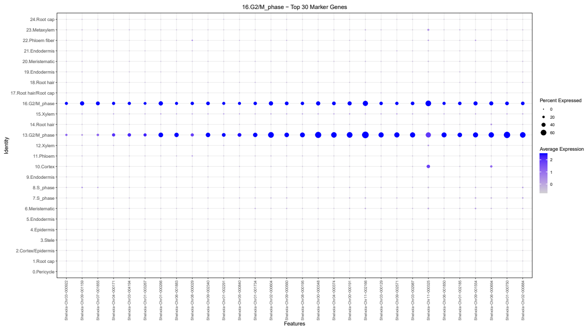
16.G2/M_phase - Top 30 Marker Genes

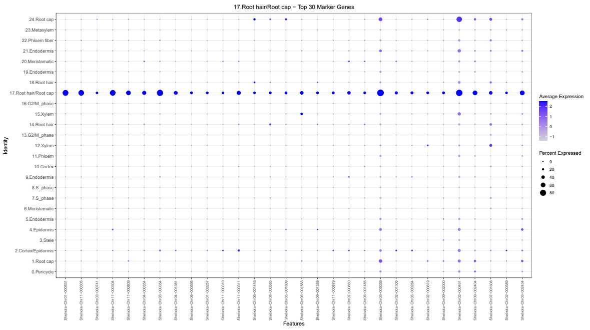
17.Root hair/Root cap - Top 30 Marker Genes

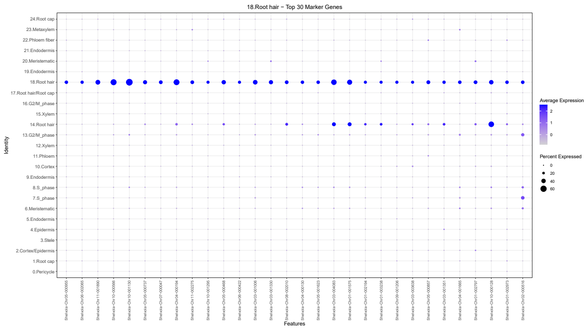
18.Root hair - Top 30 Marker Genes

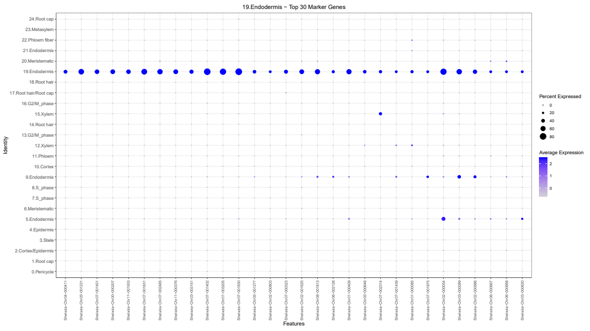
19.Endodermis - Top 30 Marker Genes

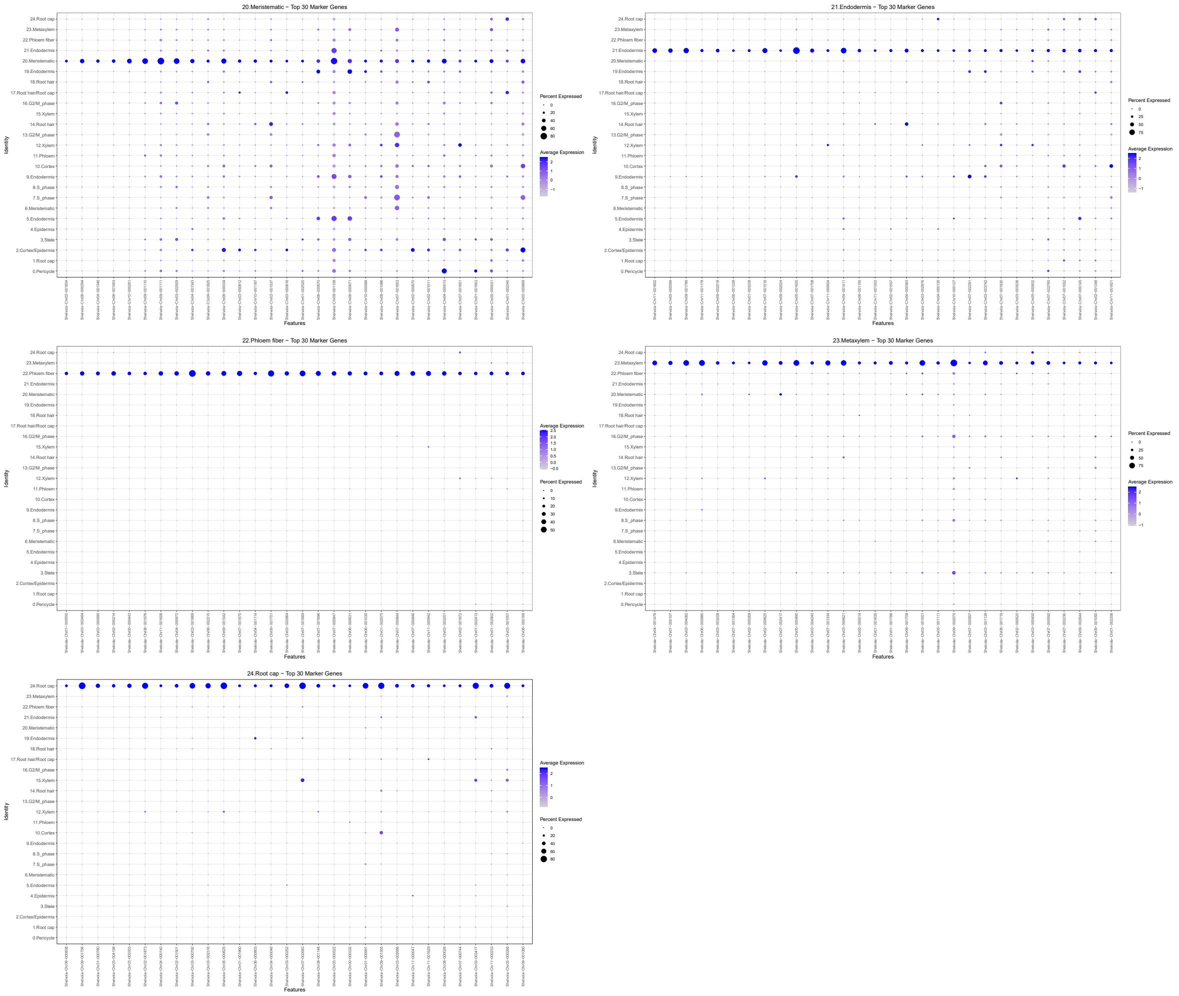


**Figure S3. Expression patterns of the top marker genes for the 25 sand bean root clusters.**

Dot plots show the top 30 positively enriched marker genes for each cluster across all 25 transcriptionally distinct clusters. Each subplot is labeled with the cluster number and corresponding cell-type annotation. Columns represent marker genes and rows represent clusters. Dot size indicates the percentage of nuclei expressing each gene, whereas dot color represents average scaled expression. The figure is displayed across three consecutive pages.

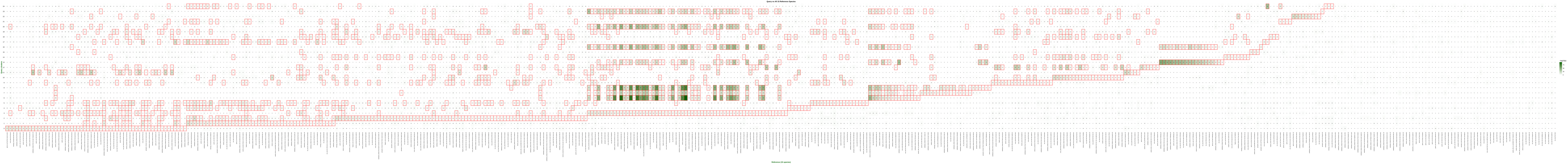

Query vs All 15 Reference Species
Reference (15 species)

**Figure S4. Cross-species OMG comparison of sand bean root clusters with reference plant cell types.**

Heatmap showing overlaps between orthologous marker gene groups from the 25 sand bean query clusters and reference cell-type marker groups compiled from 15 plant species. Rows represent sand bean clusters and columns represent reference cell types grouped by species. Cell values and shading indicate the number of shared Orthologous Marker Gene Groups (OMGs). Red outlines indicate significant cluster–reference cell-type associations identified by the orthology-guided comparison.

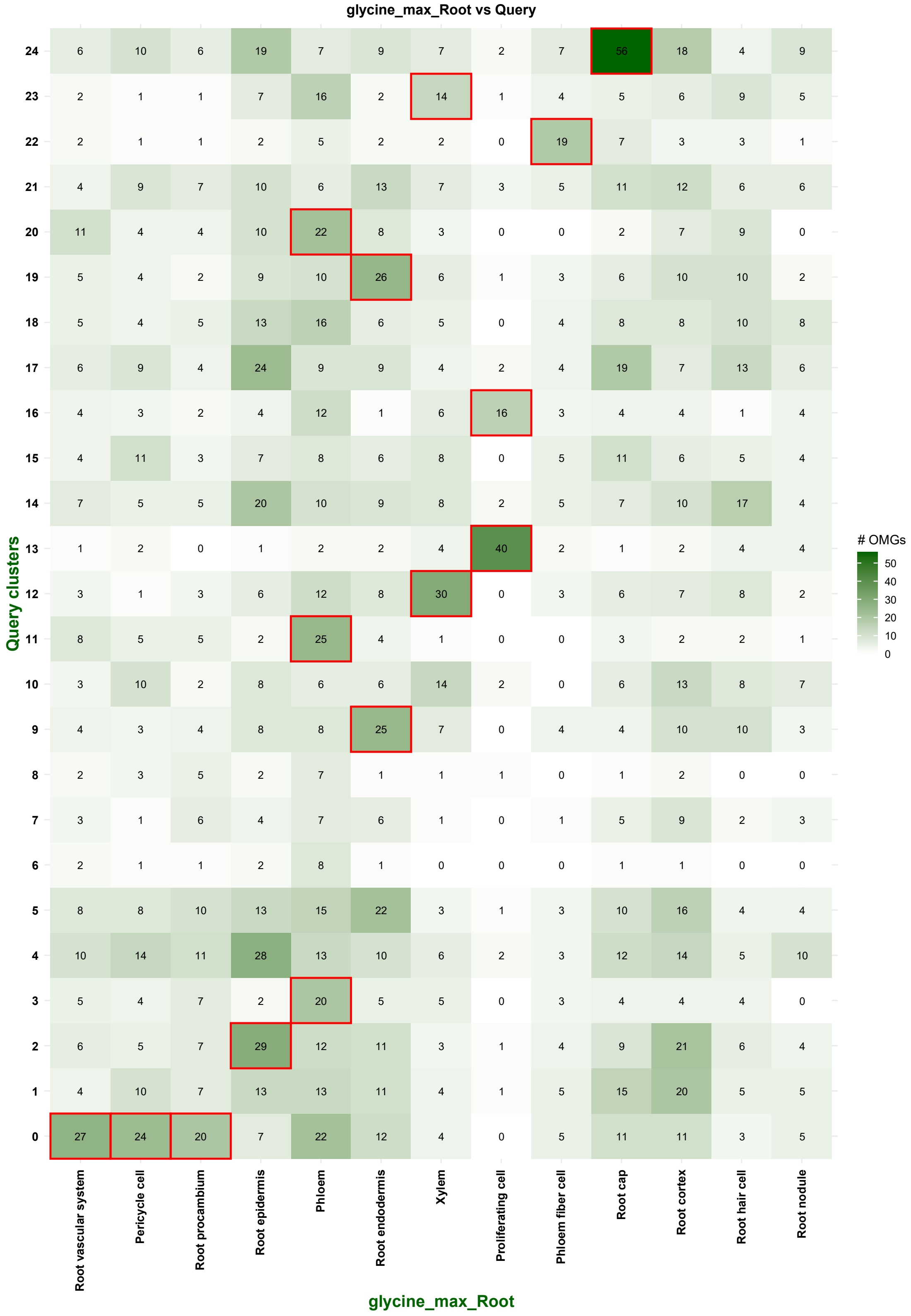


**Figure S5. OMG comparison of sand bean root clusters with soybean root cell types.**

Heatmap showing the number of OMGs shared between the 25 sand bean query clusters and reference root cell types from soybean (*Glycine max*). Rows represent sand bean clusters and columns represent soybean root cell types. Numbers and color intensity indicate the number of shared OMGs, and red outlines mark significant cluster–cell-type associations.

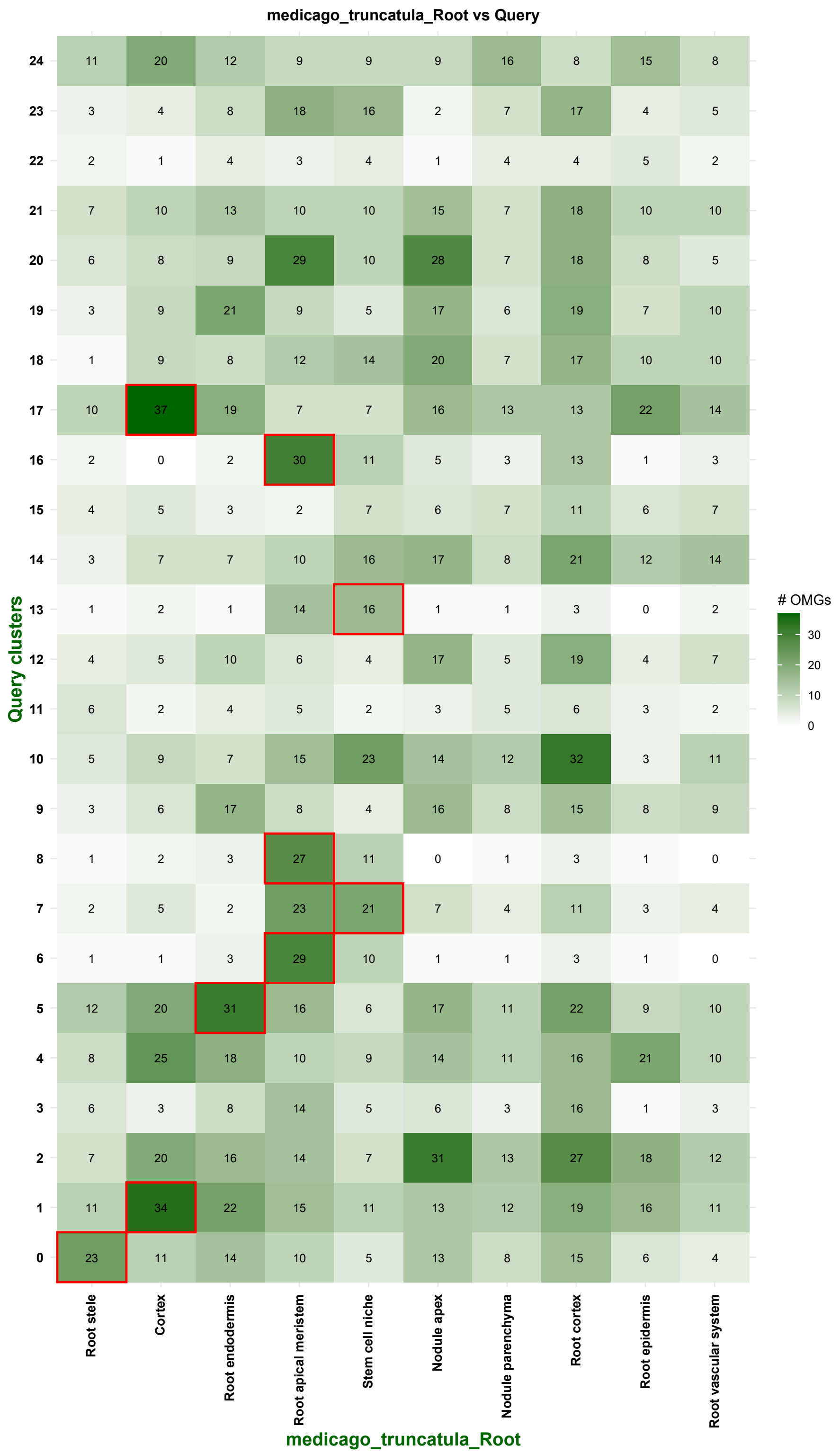


**Figure S6. OMG comparison of sand bean root clusters with *Medicago truncatula* root cell types.**

Heatmap showing OMG overlaps between the 25 sand bean query clusters and reference root cell types from *M. truncatula*. Rows represent sand bean clusters and columns represent reference cell types. Numbers and color intensity indicate the number of shared OMGs, whereas red outlines identify significant cluster–cell-type associations.

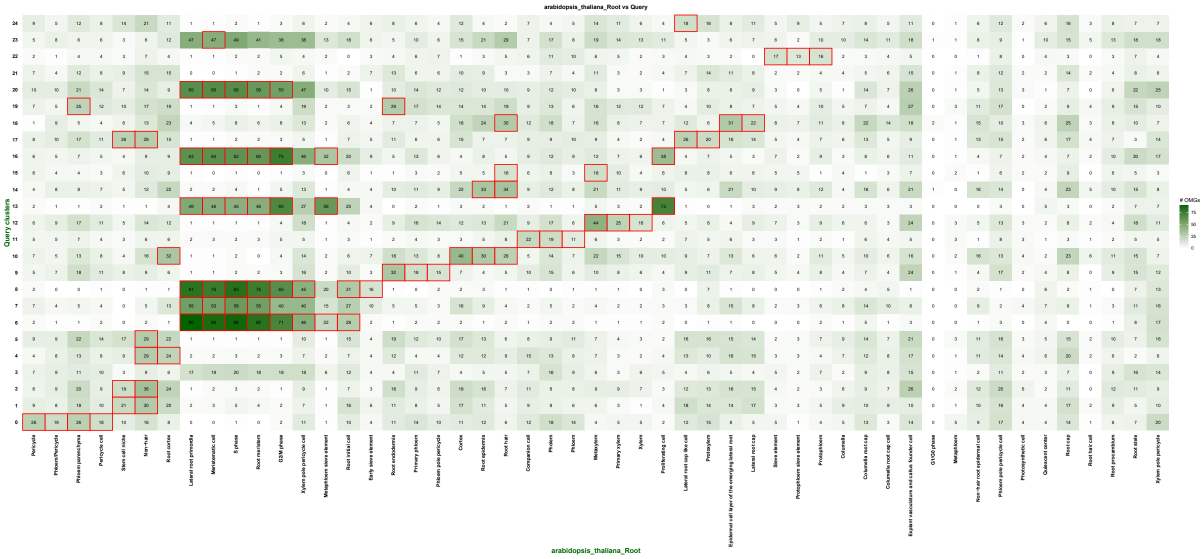


**Figure S7. Orthology-guided comparison of sand bean root clusters with *Arabidopsis thaliana* root cell types.**

Heatmap showing OMG overlaps between the 25 sand bean query clusters and reference root cell types from *A. thaliana*. Rows represent sand bean clusters and columns represent Arabidopsis root cell types. Numbers and color intensity indicate the number of shared OMGs. Red outlines indicate significant cluster–cell-type associations identified by the orthology-guided analysis.

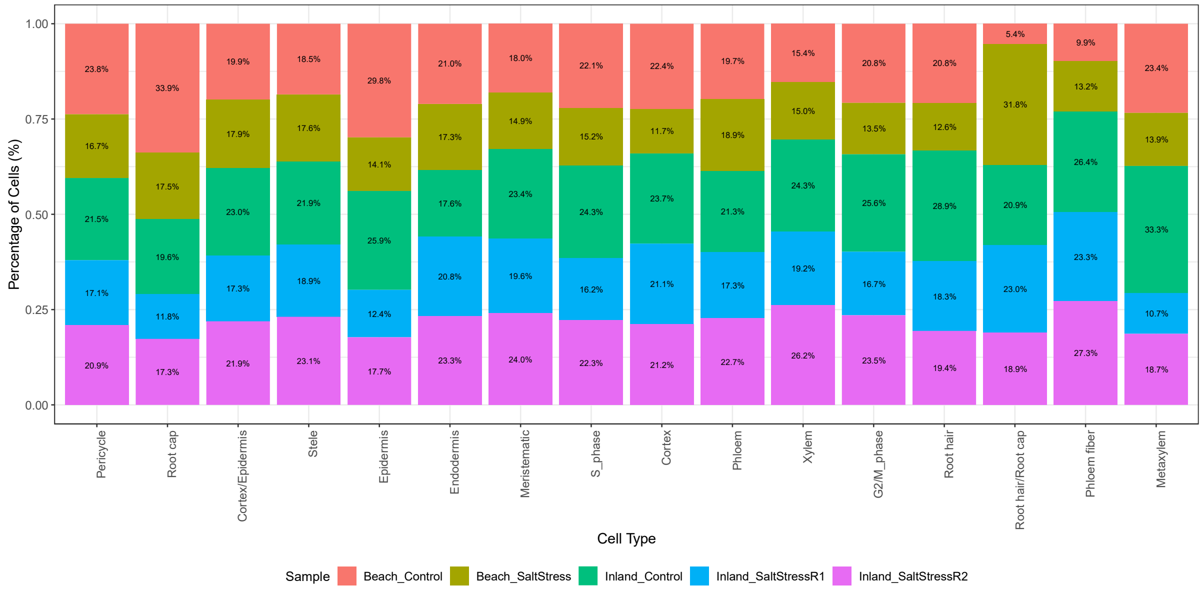

**Figure S8. Relative contributions of the five datasets to annotated sand bean root cell types.**

Stacked bar plots show the percentage contribution of Beach-Control, Beach-SaltStress, Inland-Control, Inland-SaltStressR1, and Inland-SaltStressR2 nuclei to each annotated cell type. Each bar represents one cell type and is normalized to 100%. Percentages within the colored segments indicate the contribution of each dataset to the corresponding cell population.

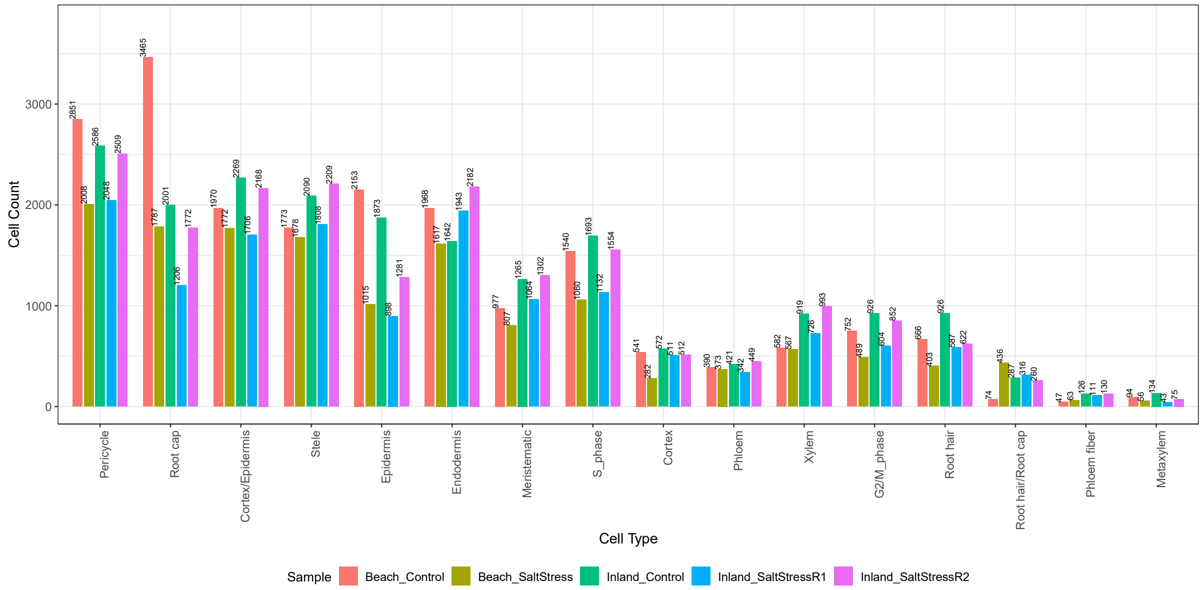

**Figure S9. Numbers of nuclei assigned to annotated root cell types across the five datasets.**

Grouped bar plots show the absolute number of nuclei assigned to each annotated cell type in Beach-Control, Beach-SaltStress, Inland-Control, Inland-SaltStressR1, and Inland-SaltStressR2. Numbers above the bars indicate the corresponding nuclei counts.

**A**

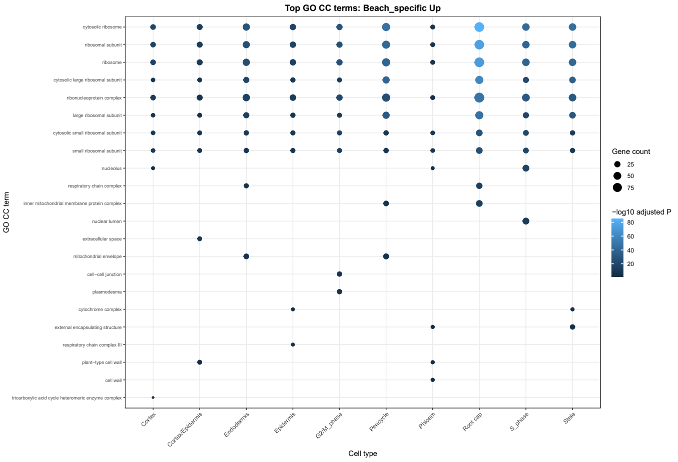


**B**

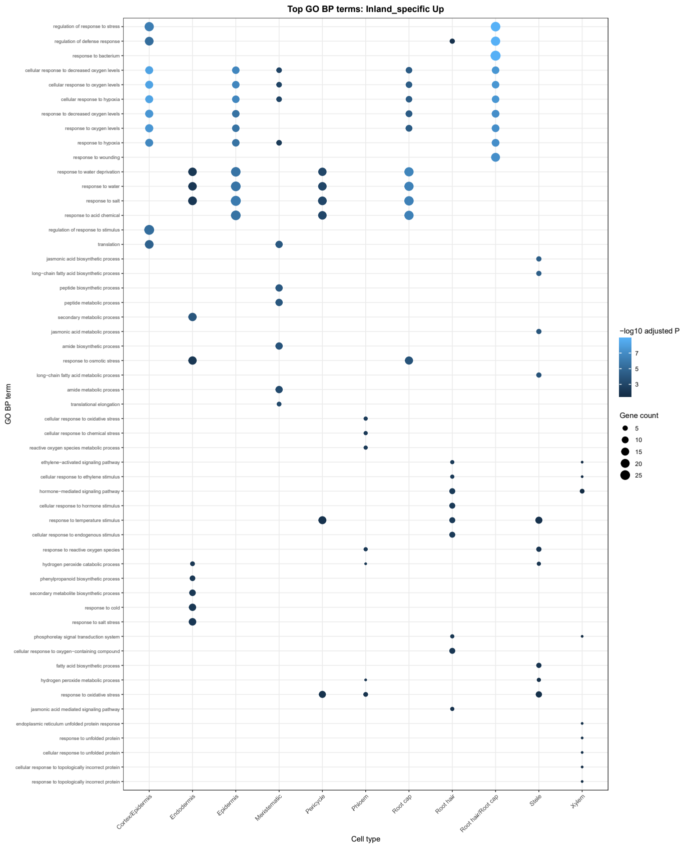


**Figure S10. Gene Ontology enrichment of ecotype-specific salt-induced genes in sand bean roots.**

**(A)** Enriched Biological Process terms among Beach-specific upregulated DEGs across individual root cell types.

**(B)** Enriched Biological Process terms among Inland-specific upregulated DEGs across individual root cell types. Columns represent cell types and rows represent enriched GO terms. Dot size indicates gene count, and dot color represents the $-\log_{10}$-transformed adjusted *P* value.
